# Sexual size dimorphism correlates with the number of androgen response in mammals, but only in small-bodied species

**DOI:** 10.1101/2024.12.07.627341

**Authors:** Caleb R. Ghione, Matthew D. Dean

## Abstract

Sexual size dimorphism (SSD) is common throughout the animal kingdom. “Rensch’s Rule” was proposed nearly 80 years ago, named for the observation that the magnitude of SSD in male-larger species increased with average body size. Here we re-examine this trend across 268 mammalian species with full genome assemblies and annotations, and place the evolution of SSD in the context of androgen response elements or estrogen response elements, the DNA motifs to which sex hormone receptors bind. Hormone receptors provide intuitive mechanisms for sex-specific regulation of the genome and could greatly impact SSD. We find that the three relatively large-bodied lineages (orders Carnivora, Cetartiodactyla, and Primates) follow Rensch’s Rule, and SSD does not correlate with the number of receptor elements. In contrast, SSD in small-bodied lineages (Chiroptera and Rodentia) correlates with the number of androgen response elements, but SSD does not correlate with overall body size. One hypothesis to unify our observations is that small-bodied organisms like bats and rodents tend to reach peak reproductive fitness quickly and are more reliant on hormonal signaling to achieve SSD over relatively short time periods. Our study uncovers a previously unappreciated relationship between SSD, body size, and hormone signaling that likely varies in ways related to life history.

## Introduction

Sexual size dimorphism (SSD) refers to the degree by which males and females of the same species differ in body size, and is a common feature across many groups including mammals ^1–3^. “Rensch’s Rule” refers to the observation that the magnitude of SSD in male-larger species tends to increase with increasing (average) body size while the magnitude of SSD in female-larger species tends to decrease. Multiple hypotheses have been proposed to account for this relationship, but all fall short of explaining why Rensch’s Rule does not seem to apply to all species.

One hypothesis to explain Rensch’s Rule is connected to life span, or at least to the age at which individuals begin reproducing. Because large-bodied species also tend to be long-lived, relatively large males will have more absolute time to accumulate body mass over multiple growing seasons. This hypothesis could also explain why small-bodied species do not follow Rensch’s Rule – because males are relatively short-lived and cannot accumulate body mass over multiple growing seasons ^4^. An intuitive corollary to this hypothesis is that small species tend to have minor SSD if the sexes have less time to accumulate different in body size.

How then do small species achieve SSD? One hypothesis is that relatively small species are more reliant upon hormonal signaling to drive males and females to different body size optima over a relatively short period of time. In other words, small-bodied species could deploy sex-specific growth rates quickly, and sex hormones (e.g., androgen and estrogen) represent a powerful molecular mechanism for sex-specific genome regulation. Androgen and estrogen circulate at different levels in males and females, and their main downstream targets are *Androgen receptor* (*Ar*) and two Estrogen receptors (*Erα* and *Erβ*). Upon binding to their hormonal ligands, these receptors undergo conformational changes, bind to particular DNA motifs, and directly or indirectly modify expression of nearby genes ^5–10^. Thus, androgen and estrogen receptors typically act as transcription factors that regulate many genes in a sex-specific manner ^11–15^, offering a potential molecular mechanism to achieve sex-specific size11,12,16-20.

Here, we test whether the magnitude of SSD correlates with the number of DNA motifs that act as androgen response elements (ARE’s) or estrogen response elements (ERE’s) near protein coding regions. We combine body size data with 268 mammalian genomes and annotations from the Zoonomia Project ^21–23^. We uncover a dramatic difference in this relationship across five orders of mammals, with the relatively small-bodied orders Chiroptera and Rodentia showing a significantly positive relationship between SSD and the number of ARE’s in their genomes. In contrast, relatively larger-bodied species from three orders (Carnivora, Cetartiodactyla, and Primates) did not show this correlation, and instead followed the classic “Rensch’s Rule”. Our study suggests that different species deploy different molecular developmental mechanisms to achieve sex-specific optima, which in turn might help resolve sexual conflict over body size.

## Results

Of 455 unique TOGA-annotated species from Zoonomia, 268 overlapped a mammalian phylogeny ^24^ and had reliable body size data in the literature (Supplementary Files 1 and 2). Among these 268 species, SSD ranged from −0.99 (dugongs) to 1.55 (southern elephant seals). Within 50 kb upstream/downstream of a transcription start site, we counted between 501 and 40,891 ARE’s (mean= 4,692) and between 12-957 (mean= 162.5) ERE’s across species.

We employed phylogenetically controlled linear models to test whether the magnitude of SSD covaried with the number of ARE’s/ERE’s and overall body size (SSD∼log(ARE-or-ERE)+log(body_size)). When all species were included, we detected a significantly positive relationship between SSD and body size (p=0.001) but not the number of ARE’s (p=0.179) or ERE’s (p=0.310) (Table 1). Closer inspection showed that these relationships varied across mammalian orders (Table 1, Figure 1, Figure 2), so we re-analyzed the five orders that were represented by at least 20 species. We could not simply include “order” as a covariate in the linear model because the scale over which SSD varies within each order is too different among groups. These five orders share almost no evolutionary history and represent hundreds of millions of years of independent evolution, justifying their separate analyses (Figure 2).

**Figure 1.**
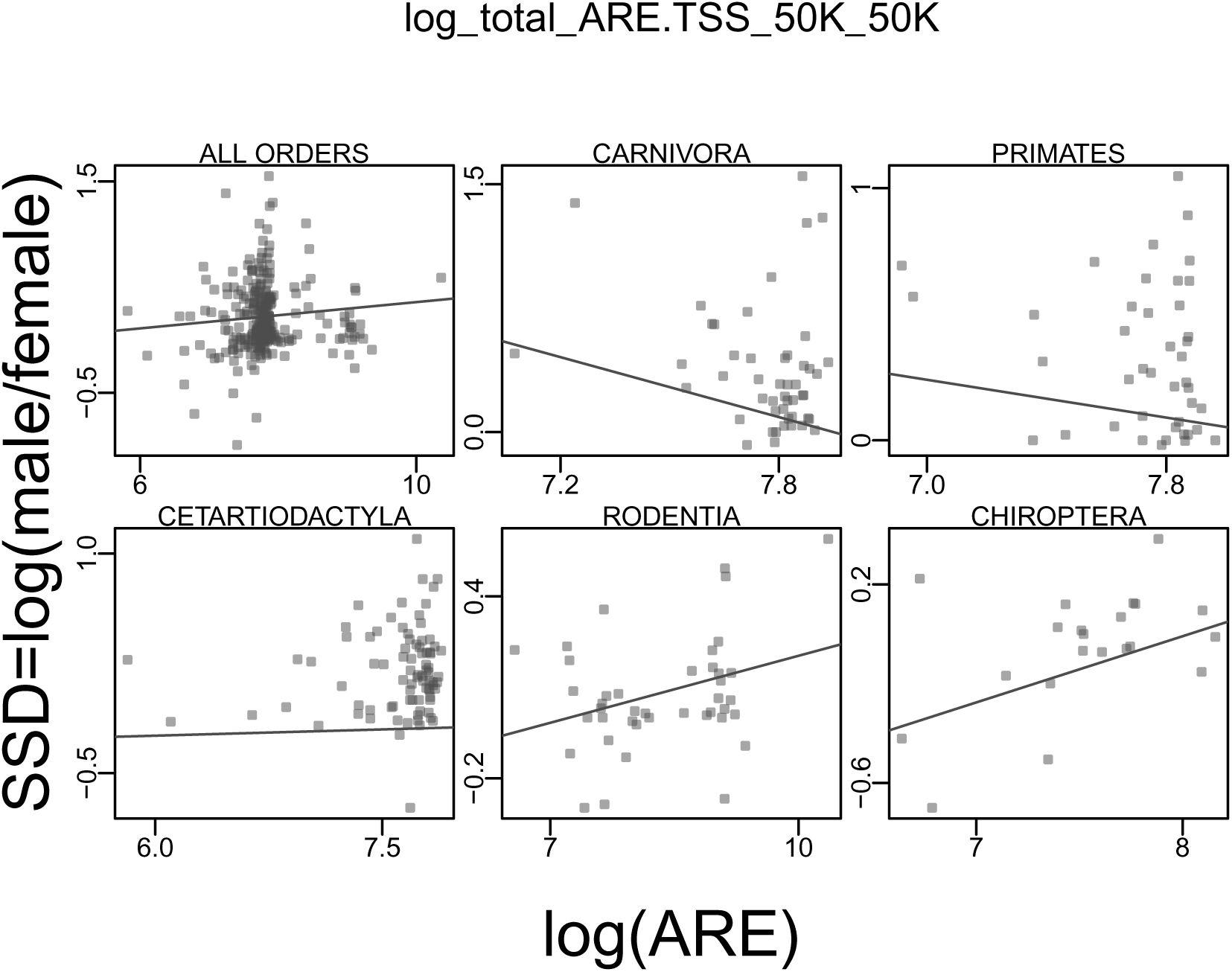
The correlations between SSD and total ARE counts among orders. ARE counts 50 kb upstream/downstream of a transcription start site, natural-log-transformed. Each point on the plot is a species, the slope of the line is derived from phylogenetically controlled linear models (see methods). The red point in Carnivora indicates the outlier species *Arctocephalus gazella* (see text).

**Figure 2.**
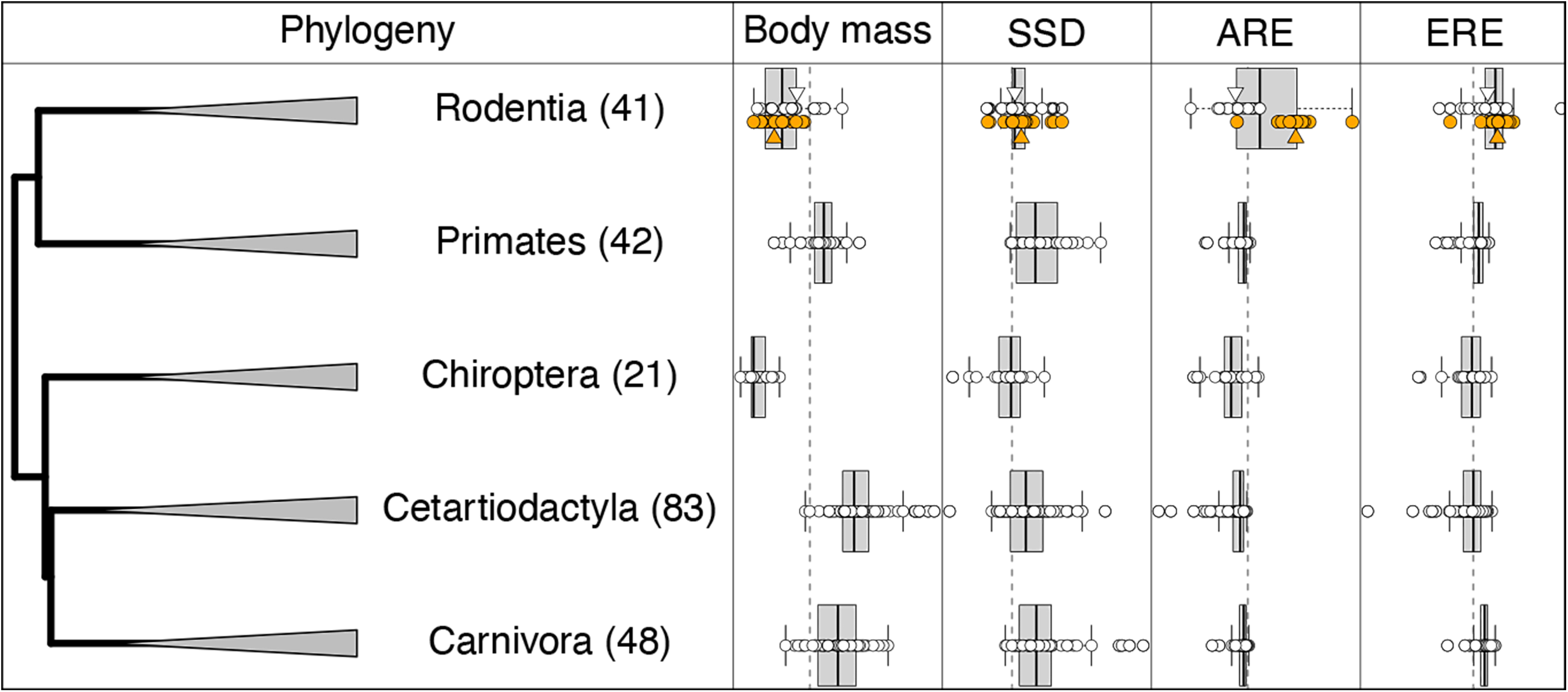
The five orders represented by at least 20 species (number in parentheses). Relationship between average body mass, SSD, ARE counts, and ERE counts. For SSD, the vertical dashed line indicates SSD=0, species to the right are male-larger, species to the left are female-larger. For all other phenotypes, the vertical dashed lines indicate phylogenetic mean estimated at the root of the mammalian tree. Rodentia has been divided into two groups for visualization: orange dots are myomorph rodents, showing their dramatic increase in ARE’s near genes. Small white and orange triangles indicate their respective means. ARE and ERE counts 50 kb upstream/downstream of a transcription start site, natural-log-transformed.

**Table 1.** phylolm results [SSD∼log(ARE or ERE) + log(bodysize)] for genomic regions within 50 Kb of transcription start site. P.values highlighted in gray indicate < 0.05.

| order | N<br>species | HRE<br>type | mean<br>HRE | HRE |  | bodysize |  | lambda | r.squared |
| --- | --- | --- | --- | --- | --- | --- | --- | --- | --- |
|  |  |  |  | estimate | p.value | estimate | p.value |  |  |
| All | 268 | ARE | 8.01 | 0.066 | 0.179 | 0.030 | 0.001 | 0.694 | 0.049 |
| Carnivora | 48 |  | 7.96 | -0.476 | 0.110 | 0.067 | 0.017 | 0.353 | 0.153 |
| Cetartiodactyla | 83 |  | 7.80 | 0.039 | 0.668 | 0.062 | 0.005 | 0.811 | 0.097 |
| Chiroptera | 21 |  | 7.74 | 0.305 | 0.034 | -0.019 | 0.747 | 0.104 | 0.228 |
| Primates | 42 |  | 7.94 | -0.221 | 0.196 | 0.064 | 0.049 | 0.437 | 0.137 |
| Rodentia | 41 |  | 8.55 | 0.078 | 0.050 | 0.006 | 0.687 | 0.000 | 0.099 |
| All | 268 | ERE | 4.72 | 0.054 | 0.310 | 0.029 | 0.001 | 0.695 | 0.046 |
| Carnivora | 48 |  | 4.84 | -0.336 | 0.265 | 0.064 | 0.043 | 0.643 | 0.099 |
| Cetartiodactyla | 83 |  | 4.54 | 0.037 | 0.663 | 0.061 | 0.005 | 0.809 | 0.096 |
| Chiroptera | 21 |  | 4.52 | 0.394 | 0.004 | -0.038 | 0.521 | 1.000 | 0.388 |
| Primates | 42 |  | 4.71 | -0.156 | 0.299 | 0.066 | 0.050 | 0.494 | 0.121 |
| Rodentia | 41 |  | 5.09 | 0.015 | 0.839 | -0.004 | 0.808 | 0.000 | 0.003 |

### In relatively large-bodied species, SSD follows Rensch’s Rule but is not correlated to ARE’s or ERE’s

Carnivora, Cetartiodactyla, and Primates all followed Rensch’s Rule, where the magnitude of SSD increased with increasing body size (p<0.017, 0.005, and 0.049, respectively) (Table 1). None of these groups showed a relationship between SSD and the number of ARE’s or ERE’s 50 kb upstream/downstream of a transcription start site. Anderson and Jones (2024) ^25^ also found no relationship between SSD and ARE’s in a study of 26 primate species. These results remain largely unchanged at if we changed the genomic window from 50 kb upstream/downstream of a transcription start site to 10 kb, 100 kb, or 1000 kb (Supplementary File 3).

### In relatively small-bodied species, SSD is positively correlated with ARE’s but does not follow Rensch’s Rule

In the two smallest-bodied lineages, Chiroptera and Rodentia, SSD was positively correlated with the number of ARE’s 50 kb upstream/downstream of a transcription start site (p=0.034 and 0.050, respectively, Table 1), but did not follow Rensch’s Rule (p=0.747 and 0.687, respectively, Table 1). These results remain largely unchanged if we changed the genomic window from within 50 kb upstream/downstream of a transcription start site to 10 kb, 100 kb, or 1000 kb (Supplementary File 3).

Interestingly, the positive correlation between SSD and ARE’s in Rodentia was accompanied by a dramatic increase in the number of ARE’s in their genomes compared to the other four orders (Figure 2). Further exploration showed this pattern was largely driven by the monophyletic clade of myomorph rodents (orange points in Figure 2). Specifically, the 21 myomorph species had nearly four times as many ARE’s on average compared to the 20 non-myomorph rodent species (13,188 vs. 3,400) (Figure 2). Myomorph rodents also had an average SSD nearly three times that of non-myomorph rodents (SSD=0.11 vs. 0.03) (Figure 2), while at the same time being an order of magnitude smaller (0.23 g vs. 3.83 g) (Figure 2).

Padilla-Morales et al. (2024) ^26^ observed an increase in the number of some gene families as a function of SSD across mammals. Therefore, we tested whether the explosion of ARE’s in myomorph rodents simply reflects an increase in the total size of the genome that was interrogated across species. This was not a likely explanation: within 50 kb upstream/downstream of a TSS, 941 Mb was interrogated from myomorph rodents on average, while 776 Mb was interrogated from non-myomorph rodents.

### Gene-centric analyses

Next, we tested each gene separately for a correlation between its ARE’s/ERE’s and SSD, again using average body size as a covariate. The order Rodentia consistently showed an excess of genes that had a positive effect on SSD. For ARE’s within 50 kb upstream/downstream of a transcription start site, 18,299 genes could be tested, meaning the gene was represented by at least 10 rodent species and had non-zero variance in ARE counts. Of these, 14,363 (78%) had a positive effect on SSD (Fig. 3). This pattern held for the 10 kb, 100 kb, and 1000 kb windows (69.0%, 82%, and 88% of genes with a positive effect on SSD, respectively). Interestingly, the opposite pattern was observed for ERE’s (Fig. 3). For the order Rodentia, 73%, 66%, and 63%, and 47% of genes (for the 10 kb, 50 kb, 100 kb, and 1000 kb cutoffs, respectively) had a *negative* effect on SSD. The pattern was less consistent among the other mammalian orders (Supplementary File 4, Supplementary File 5).

**Figure 3.**
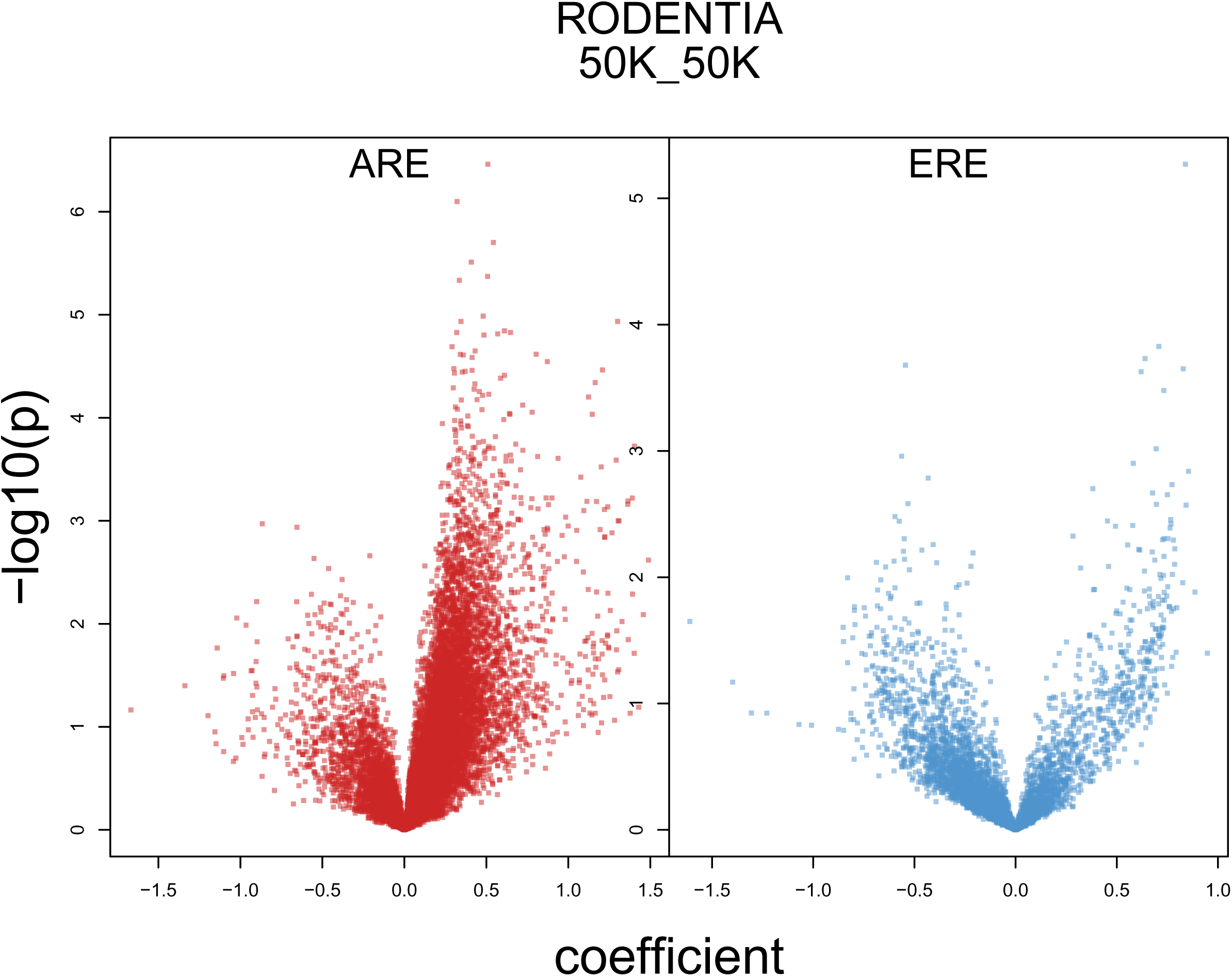
The -log10 x p-value and estimated effect size of ARE’s (red) and ERE’s (blue) per gene for Rodentia at the 50 kb upstream/downstream of TSS. For ARE’s, a large majority of genes, regardless of statistical significance, has a positive influence on SSD. This pattern contrasts with ERE’s, where the majority have negative impacts on SSD.

Given this result, we used Gene Ontology analyses to test whether genes with a significantly positive effect between their ARE’s and SSD (uncorrected p<0.05) were enriched for annotated functions. Beyond a few non-specific terms related to RNA metabolism, the Gene Ontology analyses did not yield enrichment of more intuitive functions (Supplementary File 6).

## Discussion

Nearly 80 years ago, what eventually became known as “Rensch’s Rule” posited that SSD increases with body size in male-larger species. Many hypotheses have been proposed to explain why selection might favor different optima for males and females ^27^, but it remains poorly understood *how* males and females develop different body size since they largely share the same genome. Here we combined modern genomic data with traditional phylogenetic methodology to re-examine Rensch’s Rule in the context of hormonal signaling. In small-bodied lineages, SSD correlated with ARE’s but not overall body size, suggesting the sexes rely on hormonal signaling to develop sex-specific size. In contrast, large-bodied lineages strictly followed Rensch’s Rule: their SSD increased with body size but did not correlate with the number of ARE’s or ERE’s.

Body size is correlated with numerous physiological and life history features that might give insight into the patterns we observe. Small-bodied species tend to reach sexual maturity quickly, living relatively short lives under relatively high metabolic rates ^28–37^, although there are many exceptions ^38–41^. On average, large-bodied species should therefore have more absolute time to amass differences in body size. In some cases, the larger sex takes longer to reach sexual maturity than the smaller sex ^42–47^. Such a strategy will be less available for small species if they are short-lived. Many rodent species live far less than one year – even if some of them can live for multiple years in a laboratory setting ^48^.

Our study suggests hormonal signaling is an important mechanism for small-bodied species to achieve SSD in relatively short lifespans. Since SSD increased with the number of ARE’s, one hypothesis is that males up-regulate genes that directly or indirectly increase body size. Consistent with this hypothesis, the especially small, and presumably short-lived, myomorph rodents have experienced an explosion of ARE’s in their genomes compared to all other mammalian species including other rodents. Furthermore, most genes showed a positive effect of their number of ARE’s on SSD in rodents, suggesting that most genes contribute to or are otherwise correlated with body size.

We currently lack the necessary data or methods to directly link hormonal signaling to SSD across these species, and the proxy we use (the numbers of ARE’s and ERE’s) potentially suffers from several limitations. First and foremost, the presence of a DNA motif does not necessarily mean it serves as a binding site for AR or ER. The 18 different ARE’s have variable binding specificity to AR and can be bound by transcription factors other than AR ^9,49–54^, and ARE’s sometimes act cooperatively to induce gene expression ^10^, obscuring simple connections between the number ARE motifs and transcription of nearby genes. Chromatin structure will also impact the accessibility of hormone binding elements to their receptors, so even if an ARE motif could impact gene expression, it might have little effect if buried in closed chromatin. Even if we knew which hormone receptors bound which elements, we would not know how gene expression is affected without further experimental manipulation, let alone how that variation contributes to variation in sex-specific body size. Perhaps such complexities explain why the relationship between androgen signaling and body size is sometimes paradoxical – for example, in two closely related species of lizards, testosterone had opposite effects on male body size ^55,56^. Nevertheless, Anderson and Jones (2024) ^25^ showed that ARE’s congregated near male-biased genes and androgen-responsive genes, supporting the assumption that hormone binding elements serve as a meaningful proxy for genomic regulation, at least with respect to AR. We would also argue that all the uncertainty of using binding motifs as a proxy for genomic regulation should only add statistical noise to the relationship with SSD, and it is possible that the true correlations are much stronger.

Because males and females largely share the same genome, genetic variation that influences body size is likely to evolve under sexual conflict, where an allele that increases body size benefits the larger sex to the detriment of the other. Our study explores mechanisms by which sexual conflict could be resolved, revealing different strategies that could be deployed across species. Our study suggests that small species rely more heavily on hormonal signaling to achieve SSD, while large species do not. Multiple studies have demonstrated sexually antagonistic genetic variation in many different species ^57–64^, suggesting that conflict is never fully resolved, however. The existence of sexually antagonistic genetic variation likely poses constraints on the magnitude of SSD ^65^, leading Pennell et al. (2013) ^58^ to coin the phrase “you can’t always get what you want”.

It is important to recognize that the mammalian lineages studied here differ not only in body size but also in the magnitude of their SSD. The three large-bodied lineages all have relatively large SSD, ranging from an average of 0.195 (Cetartiodactyla) to 0.375 (Carnivora). In contrast, the small-bodied lineages have fairly modest SSD of 0.073 (Rodentia) and in fact shows female-larger SSD of −0.058 in Chiroptera, a pattern consistent with other studies ^4,66^. Rensch’s Rule applies to male-larger species, so another explanation is that SSD in smaller-bodied lineages is too small. However, the positive correlation between SSD and ARE’s in both rodents and bats, the explosion of ARE’s in the especially small myomorph rodents, and the pervasive positive effect on SSD across genes, all suggest that small-bodied lineages are selected to achieve SSD, even if modest, through hormonal signaling.

Our study unites the classic Rensch’s Rule with the annotation of hundreds of mammalian genomes. The magnitude of SSD varies with the number of ARE’s, but only in small-bodied lineages. Large-bodied species strictly follow Rensch’s Rule. Our study suggests that hormonal signaling is an important mechanism enabling males and females to approach different body size optima in small-bodied species. Future studies linking molecular biology of androgen signaling and body size promise to shed light on the fundamentally important evolution of body size in males vs. females, and its potential relation to sexual conflict.

## Methods

### Body Mass

We collected body mass data taken outside of breeding season, primarily to avoid data from pregnant or lactating females. Most body mass data were taken from Silva and Downing (1995) ^67^, supplemented with additional literature sources (Supplementary File 1) ^67–75^. Sexual size dimorphism (SSD) was calculated as *ln*(male/female) body mass.

### ARE and ERE counts across whole genomes

We counted ARE’s and ERE’s across all genome assemblies of the Zoonomia consortium, using genome annotations from the Zoonomia Project ^21–23^. Zoonomia sequenced and assembled (or linked to genomes stored in NCBI) 524 genome assemblies from 464 unique species.

The canonical ARE is AGAACANNNTGTTCT, where the three N’s indicate a three nucleotide spacer flanked by palindromic sequence ^49,52,76–78^. 17 additional ARE’s have been identified, all of which are similar in sequence but vary in their specificity of binding with AR. Following Anderson and Jones (2024) ^25^, we analyzed all 18 ARE’s pooled. A single estrogen receptor element (ERE) has been identified as the sequence GGTCANNNTGACC ^5,8,78–80^. We counted ARE’s and ERE’s using the VCOUNTPATTERN function from the BIOSTRINGS package in R (v 2.68.1) ^81^. For any non-palindromic ARE’s, we added up the counts in the forward and reverse-complement directions of the genome.

### ARE and ERE counts near protein coding regions

To focus on protein coding regions and their *cis*-regulatory regions, we used the gene annotations of Kirilenko et al. (2023) ^82^, who developed *A Tool to infer Orthologs from Genome Alignments* (TOGA) to essentially lift over annotations from a reference genome (the house mouse genome annotation *mm10*) to all other genome assemblies. TOGA makes six different gene calls: i) *Intact* genes, where the middle 80% of the *mm10* reference gene is identified without inactivating mutations (frameshifts or early termination) in the target genome, ii) *Partially intact* genes, which are intact genes that show gaps in the genome assembly, iii) *Lost* genes, where the middle 80% of the *mm10* gene is present but contains at least one inactivating mutation, iv) *Missing* genes, where less than 50% of the *mm10* gene is present, v) *Uncertain lost* genes, which are genes that would be classified as *Lost* but where evidence is not strong, and vi) *Paralogous projection* genes, which are *Intact* genes that are not orthologous to the reference *mm10* gene, for example a lineage-specific duplication of a gene.

We counted ARE’s and ERE’s that fell 10 Kb, 50 Kb, 100 Kb, or 1000 Kb upstream/downstream of transcription start sites of either *Intact* or *Paralogous Projection* genes. We included *Paralogous Projection* genes because our hypothesis was focused on sex-specific expression of the genome, regardless of whether a gene was duplicated. We counted ARE’s and ERE’s once, even if they fell near multiple protein coding genes. The proximity over which ARE’s and ERE’s can affect gene expression is a topic of debate, but ARE’s have been shown to influence genes on the order of 1000 Kb away ^83^. We arrive at highly similar conclusions regardless of which cutoff we use, but present results based on the 50 Kb cutoff in the main manuscript and discuss differences when they arise.

### Phylogenetically controlled linear modeling

We implemented phylogenetically controlled linear models using the PHYLOLM function in the R package PHYLOLM ^84^ The basic model tested was SSD∼ARE (or ERE) counts + body_size. SSD was calculated as *ln*(male/female body mass). ARE (or ERE) counts were counts from each genome. Body size was calculated as of the average male+female body mass in each species. ARE (or ERE) counts and body size were *ln*-transformed prior to analyses. Average body size was included as a covariate because of “Rensch’s Rule”, the observation that SSD in male-larger species increases with body size ^85^. In addition, many life history traits covary with body size. We used the phylogeny of Upham et al. ^24^, trimmed to match our data (Supplemental File 2).

### Gene-centric analyses

We also tested whether SSD correlated with ARE’s or ERE’s on a gene-by-gene basis. This analysis represents an agnostic approach toward revealing genes or gene classes that explained variation in sexual size dimorphism. As above, we counted ARE’s and ERE’s that fell within 10 Kb, 50 Kb, 100 Kb, and 1000 Kb upstream or downstream of the transcription start site of *Intact* and *Paralogous Projection* genes. In this gene-centric approach only, ARE’s/ERE’s could be counted more than once if they fell near multiple genes. We only analyzed genes that were represented by at least 10 species and showed non-zero variance in ARE or ERE counts.

### Gene ontology analysis

Following our gene-centric analyses, we tested whether individual genes that were significantly positively correlated with SSD (uncorrected p<0.05) were over-represented for Gene Ontology terms, using PantherDB ^86^.

### Genome quality

Some of our analyses could be confounded by variation in genome quality. For example, Kirilenko et al. (2023) ^82^ annotated more genes from relatively high-quality genomes. This should not be a significant problem in the current study since we confined our analyses to protein coding regions, which are generally easier to assemble than intergenic regions which often contain large repetitive elements. Nevertheless, we tested whether ARE or ERE counts correlated with contig N50, a common metric of genome quality, taken from Kirilenko et al. (2023) ^82^. N50 is the contig length or longer which includes half the bases of the assembly.

## Supporting information

SuppFile1

SuppFile2

SuppFile3

SuppFile4

SuppFile5

SuppFile6

## Acknowledgements

Diane Genereux (Broad Institute) fielded many inquiries regarding Zoonomia data downloads; without her help this project would have been impossible. We thank Drew Anderson for many productive discussions on sexual size dimorphism, and Matt Pennell for many useful discussions on phylogenetic methodology.

---

**Supplemental File 1.** All species, ARE’s, and ERE’s analyzed in the manuscript.

**Supplemental File 2.** The phylogenetic tree relating the species analyzed in the manuscript.

**Supplemental File 3.** All phylogenetically controlled linear models performed, across all subsets of data.

**Supplemental File 4.** A repeat of Figure 3, but for all other orders and genomic window cutoffs.

**Supplemental File 5.** The proportion of gene that show a positive or negative correlation between the number of their ARE’s or ERE’s and SSD, including all orders and genomic window cutoffs.

**Supplemental File 6.** Results from Gene Ontology analyses, only for genes that whose number of ARE’s were significantly (uncorrected p<0.05) and positively correlated to SSD, only for Rodentia and the 50 kb upstream/downstream of TSS.

## Notes

### Competing Interest Statement

The authors have declared no competing interest.

