## Supplementary material for "Sexual size dimorphism correlates with the number of androgen response in mammals, but only in small-bodied species": SuppFile4

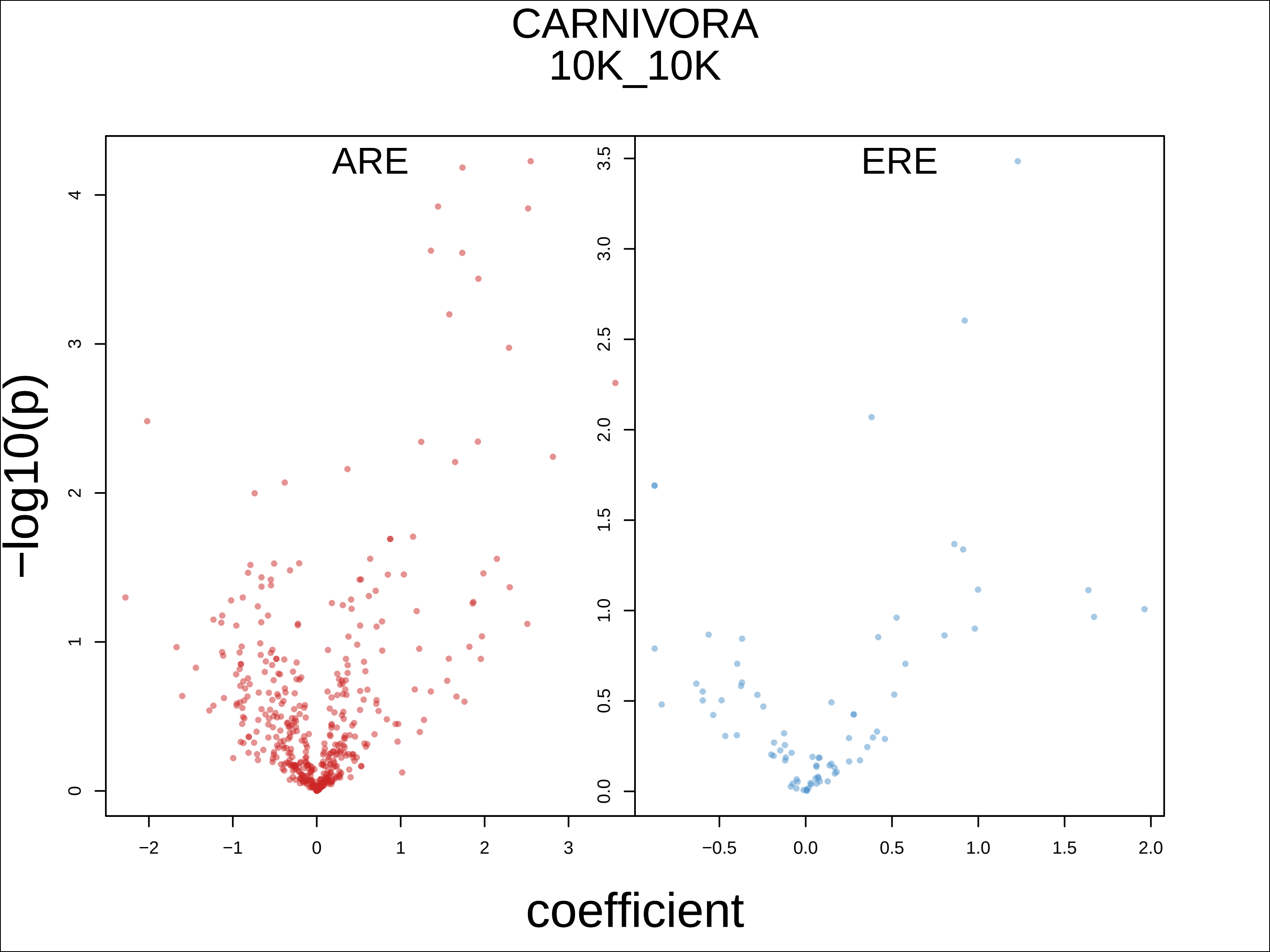

CARNIVORA
10K_10K
ARE
ERE
-log10(p)
coefficient
0
1
2
3
4
−2
−1
0
1
2
3
0.0
0.5
1.0
1.5
2.0
2.5
3.0
3.5
−0.5
0.0
0.5
1.0
1.5
2.0

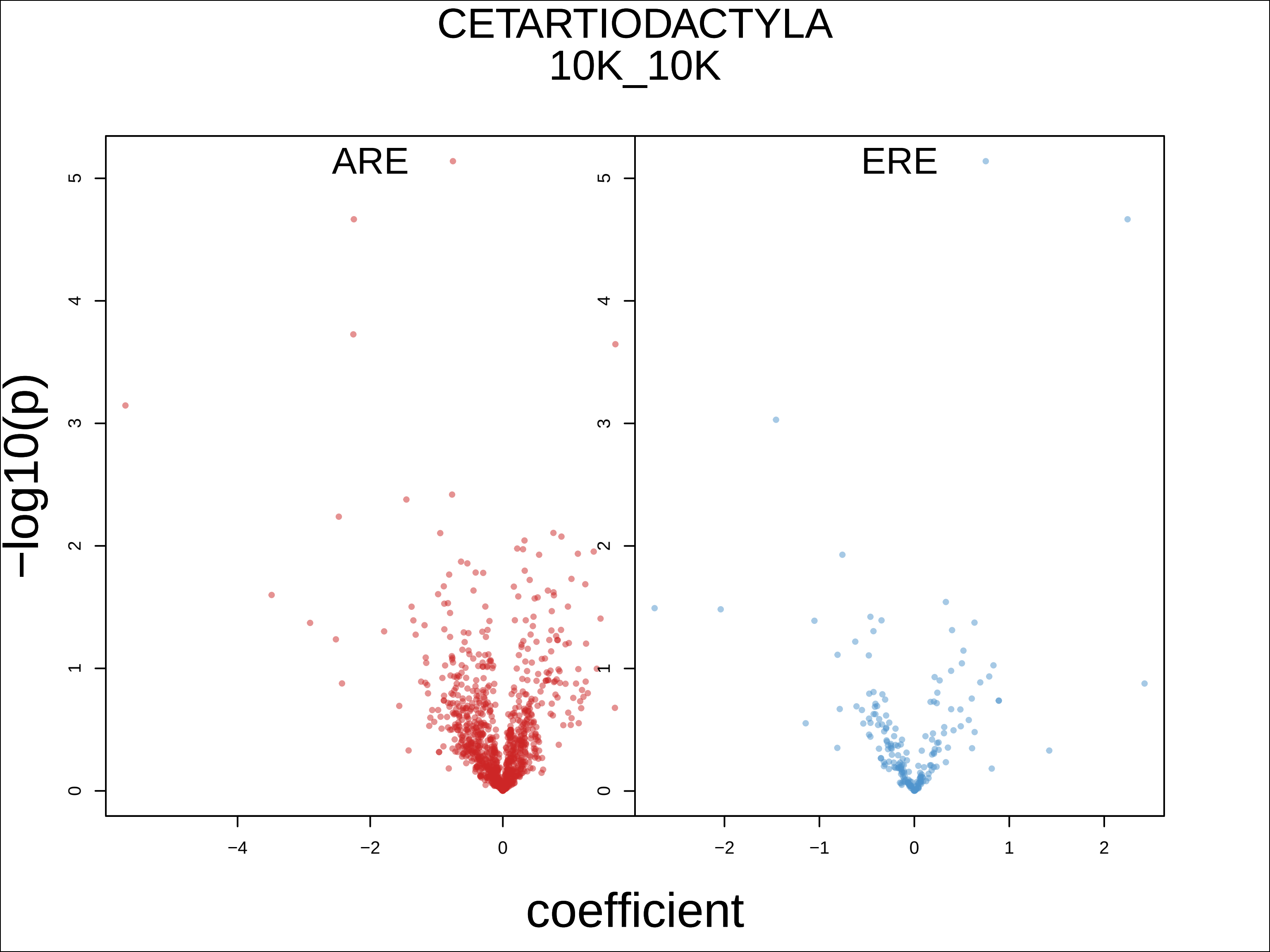

CETARTIODACTYLA
10K_10K
ARE
ERE
−log10(p)
coefficient
0
1
2
3
4
5
−4
−2
0
−2
−1
0
1
2

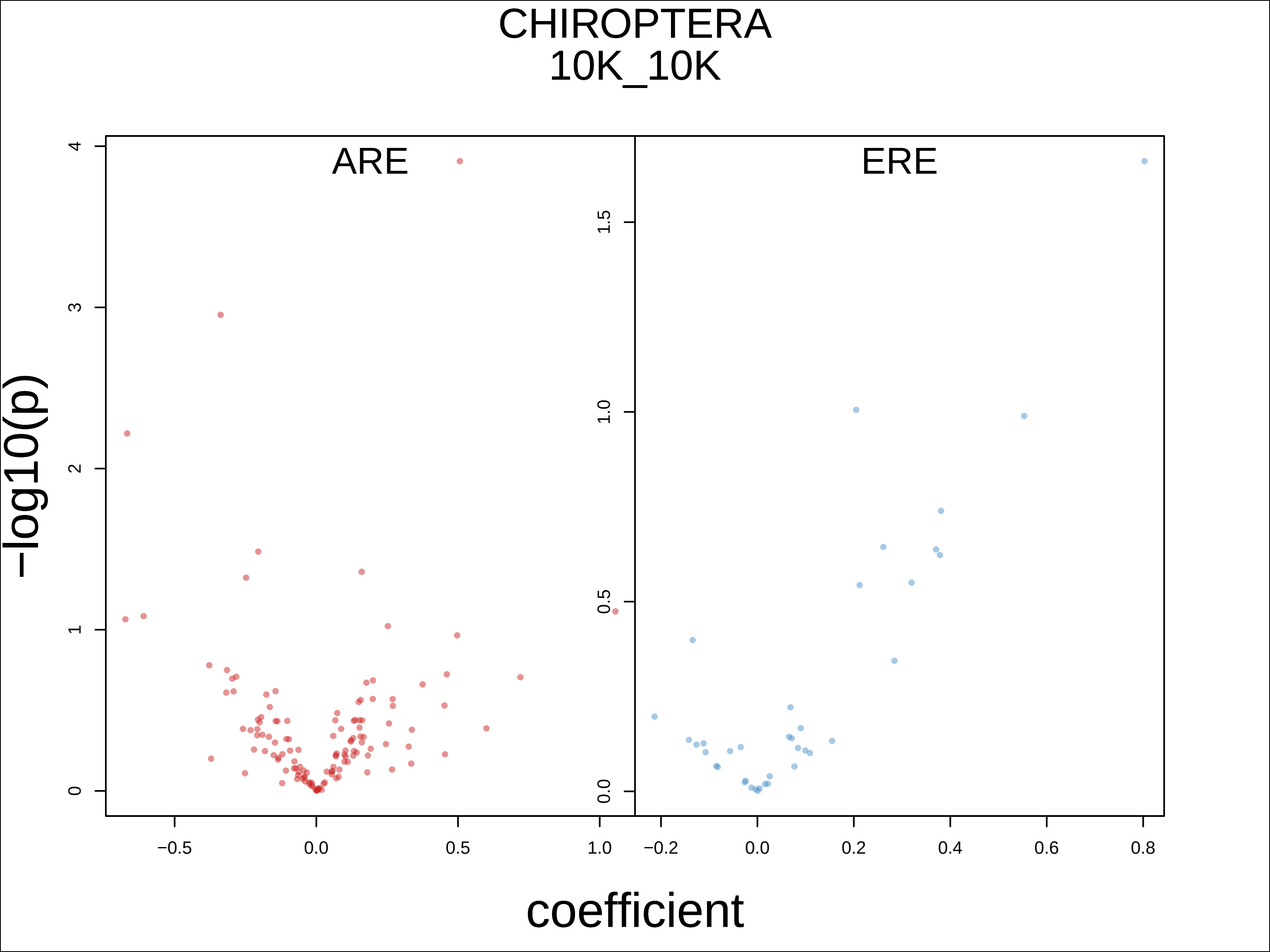

CHIROPTERA
10K_10K
ARE
ERE
−log10(p)
coefficient
0
1
2
3
4
0.0
0.5
1.0
1.5
−0.5
0.0
0.5
1.0
−0.2
0.0
0.2
0.4
0.6
0.8

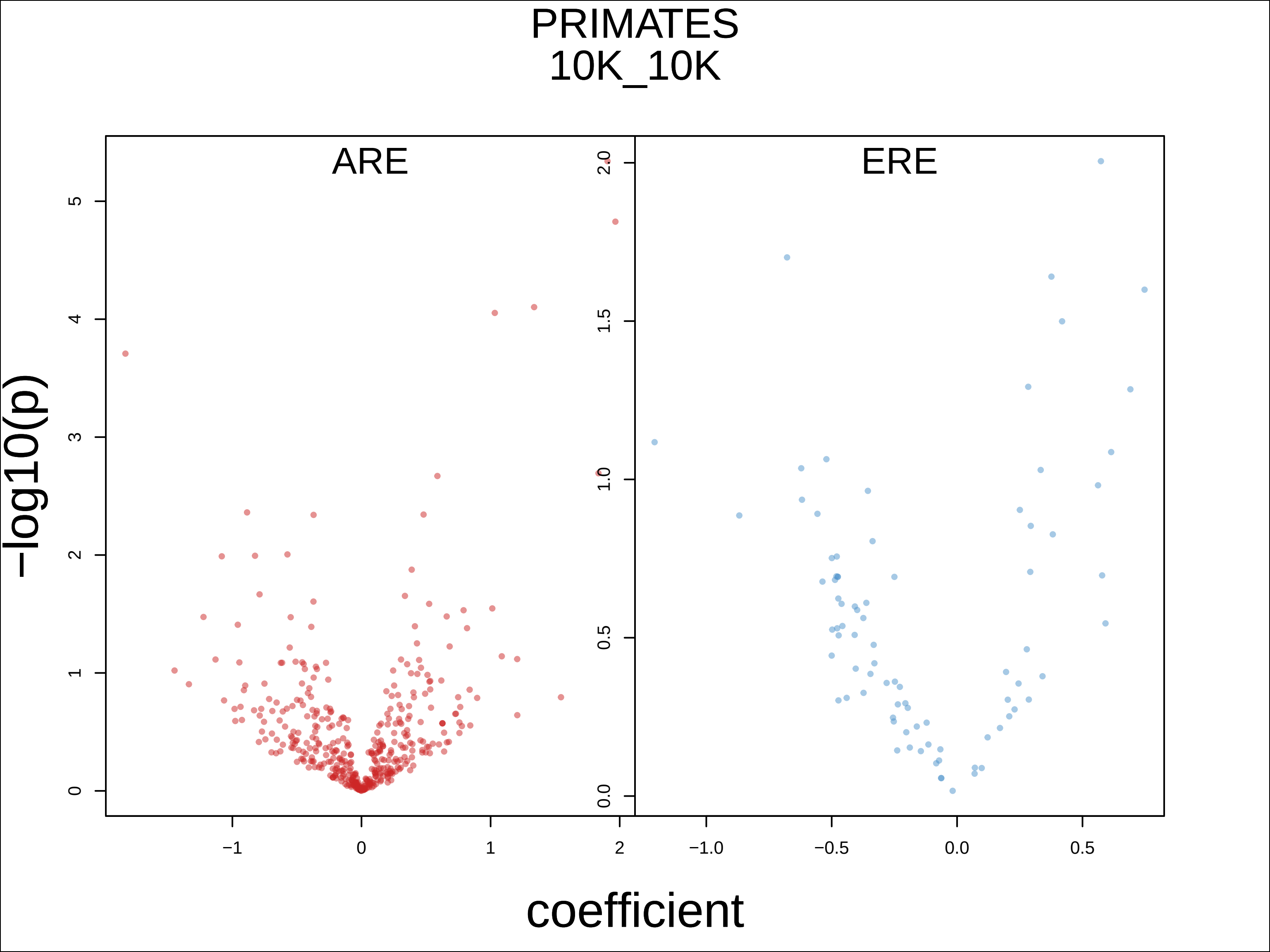
PRIMATES
10K_10K
ARE
ERE
−log10(p)
coefficient
0
1
2
3
4
5
−1
0
1
2
0.0
0.5
1.0
1.5
2.0
−1.0
−0.5
0.0
0.5

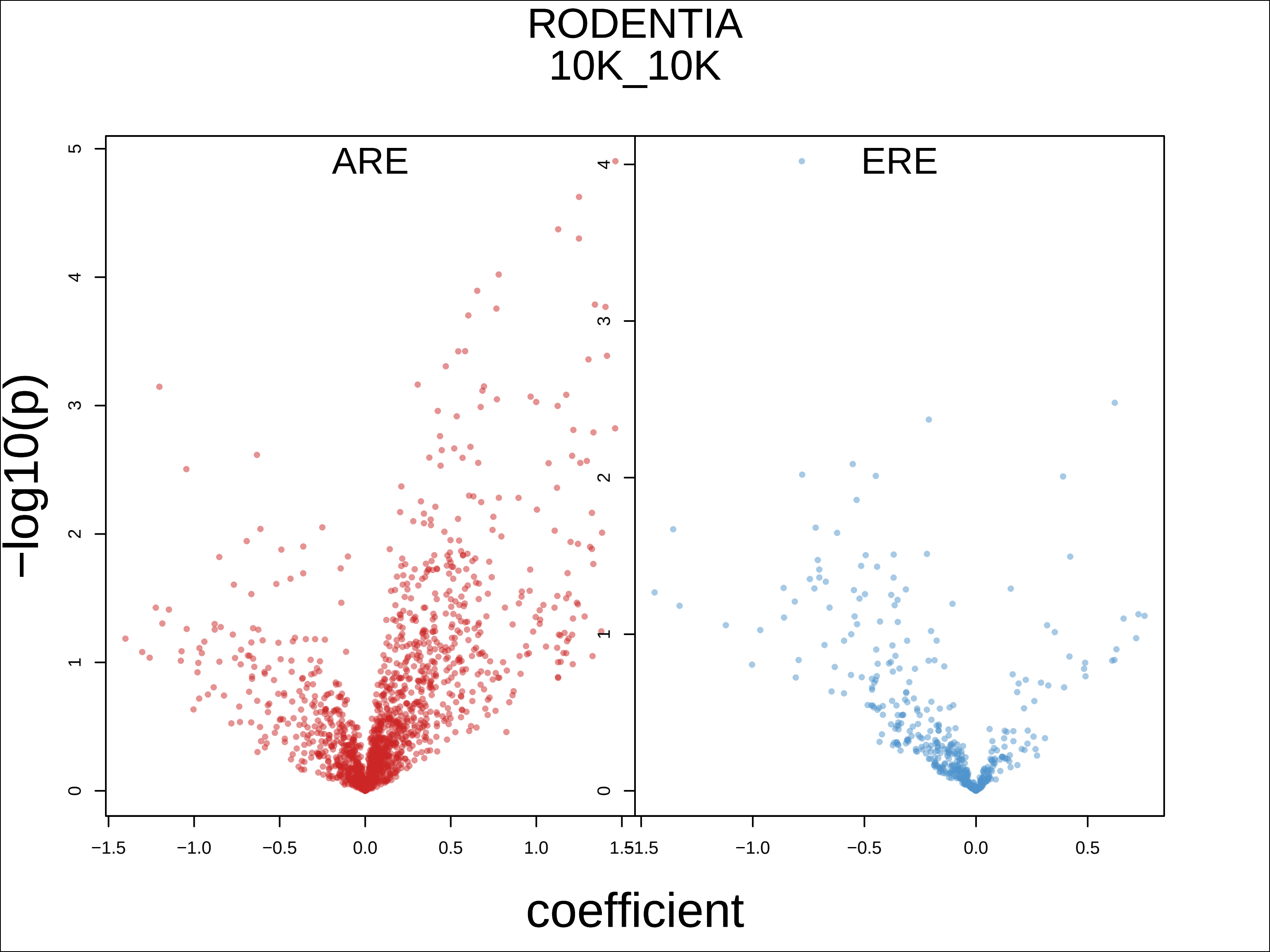

RODENTIA
10K_10K
ARE
ERE
−log10(p)
coefficient
0
1
2
3
4
5
−1.5
−1.0
−0.5
0.0
0.5
1.0
1.5

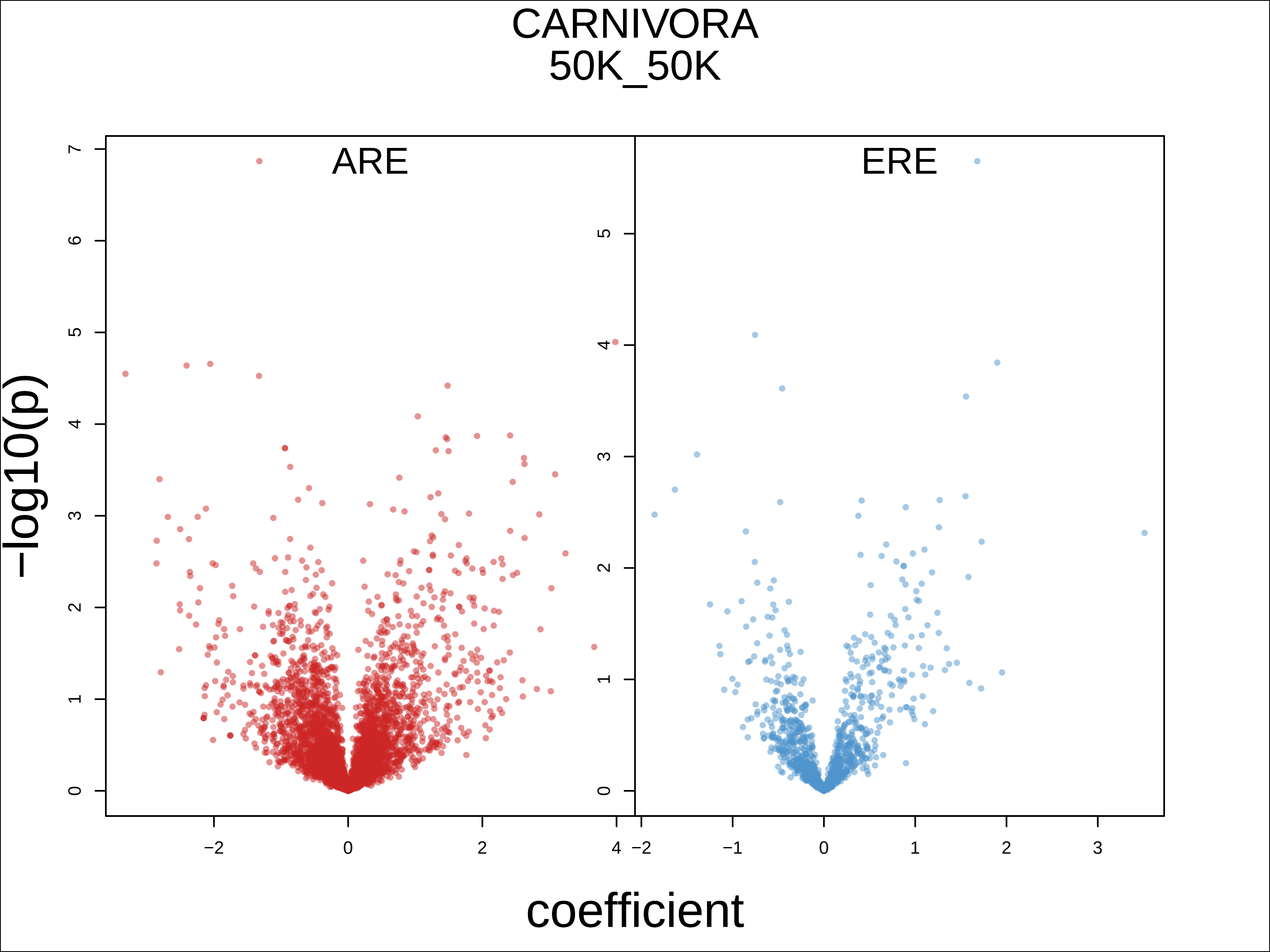
CARNIVORA
50K_50K
ARE
ERE
−log10(p)
coefficient
0
1
2
3
4
5
6
7
−2
−1
0
1
2
3
4

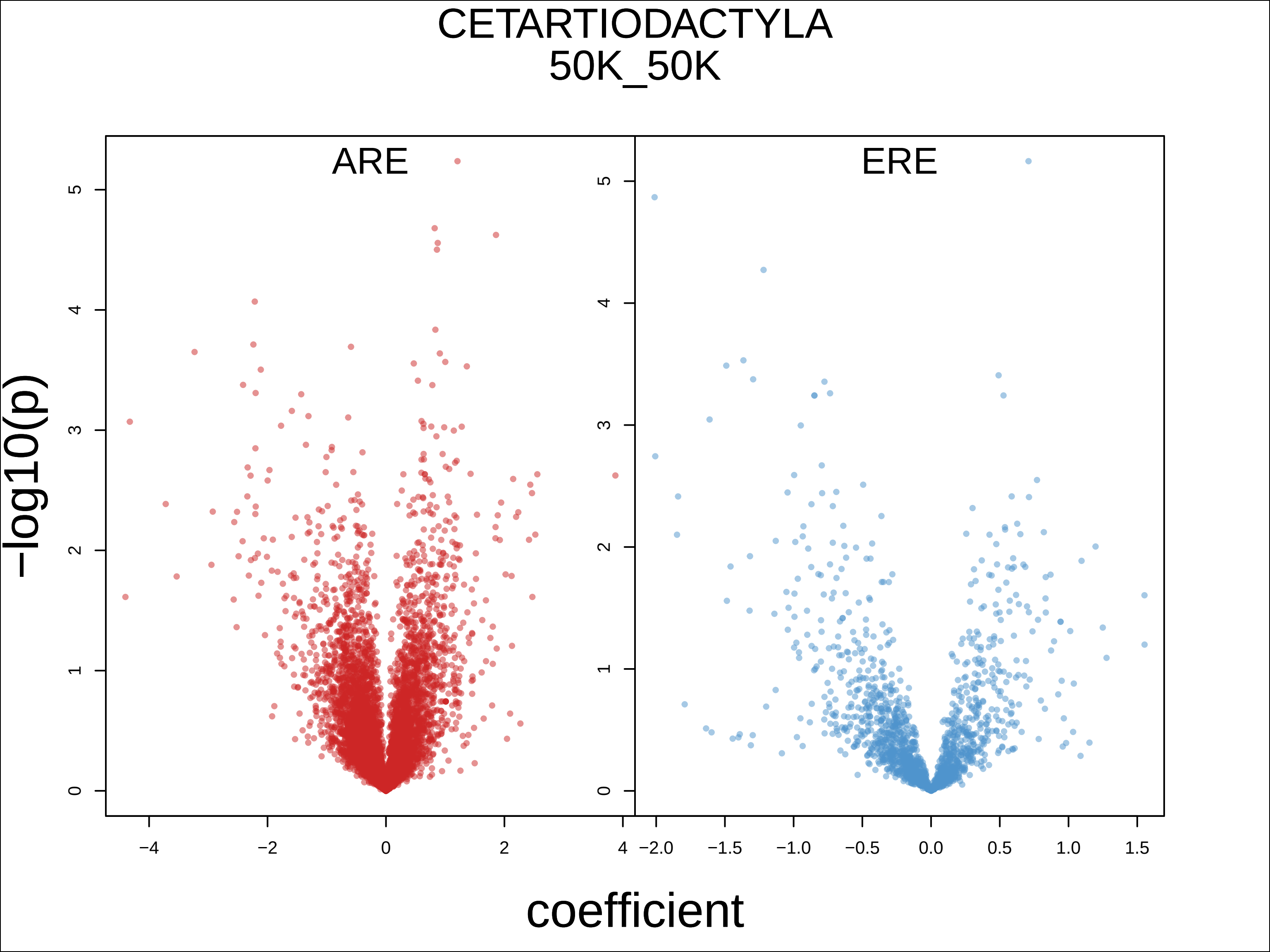

CETARTIODACTYLA
50K_50K
ARE
ERE
−log10(p)
coefficient
0
1
2
3
4
5
−4
−2
0
2
4
−2.0
−1.5
−1.0
−0.5
0.0
0.5
1.0
1.5

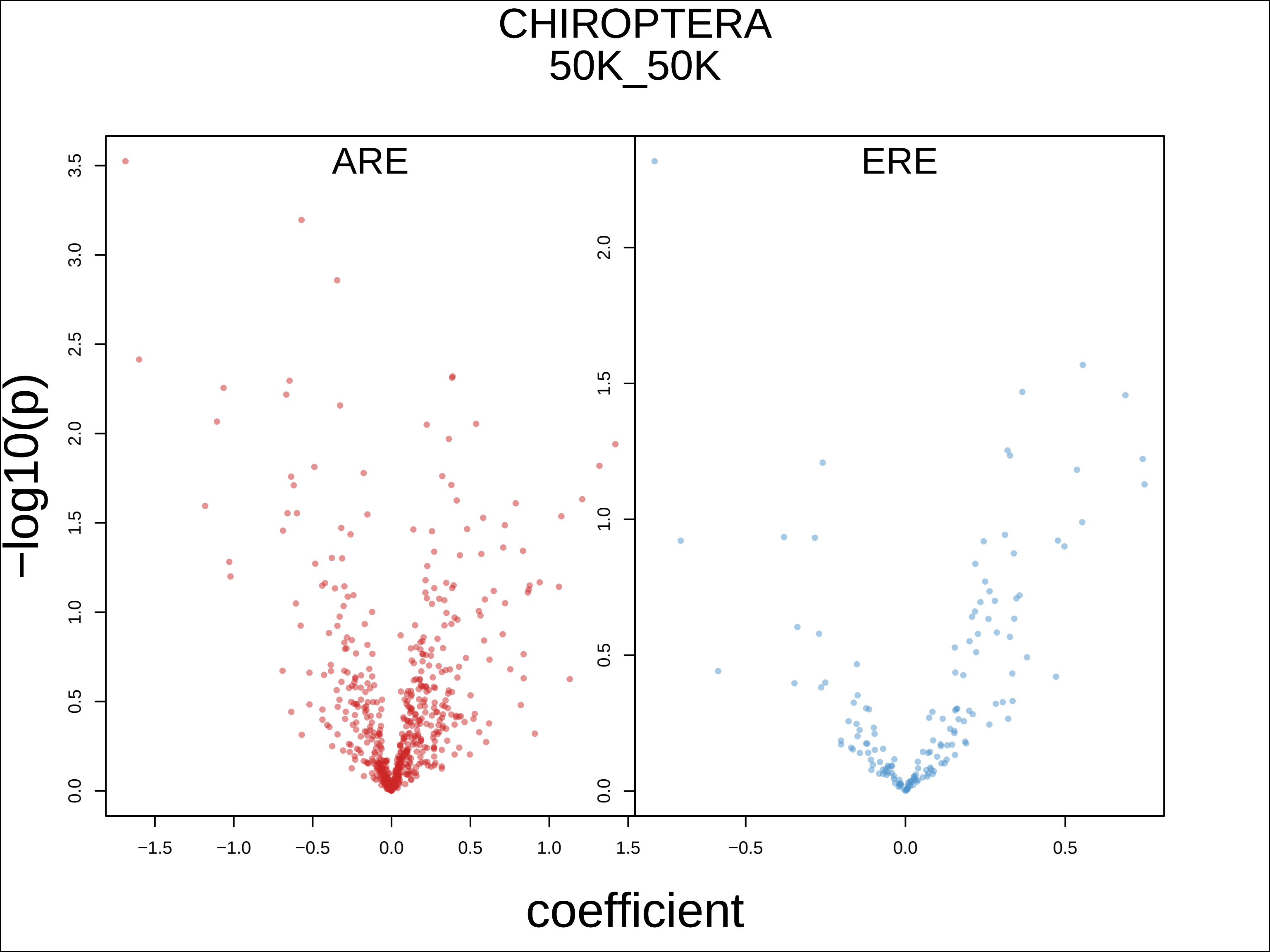

CHIROPTERA
50K_50K
ARE
ERE
−log10(p)
coefficient
0.0
0.5
1.0
1.5
2.0
2.5
3.0
3.5
−1.5
−1.0
−0.5
0.0
0.5
1.0
1.5

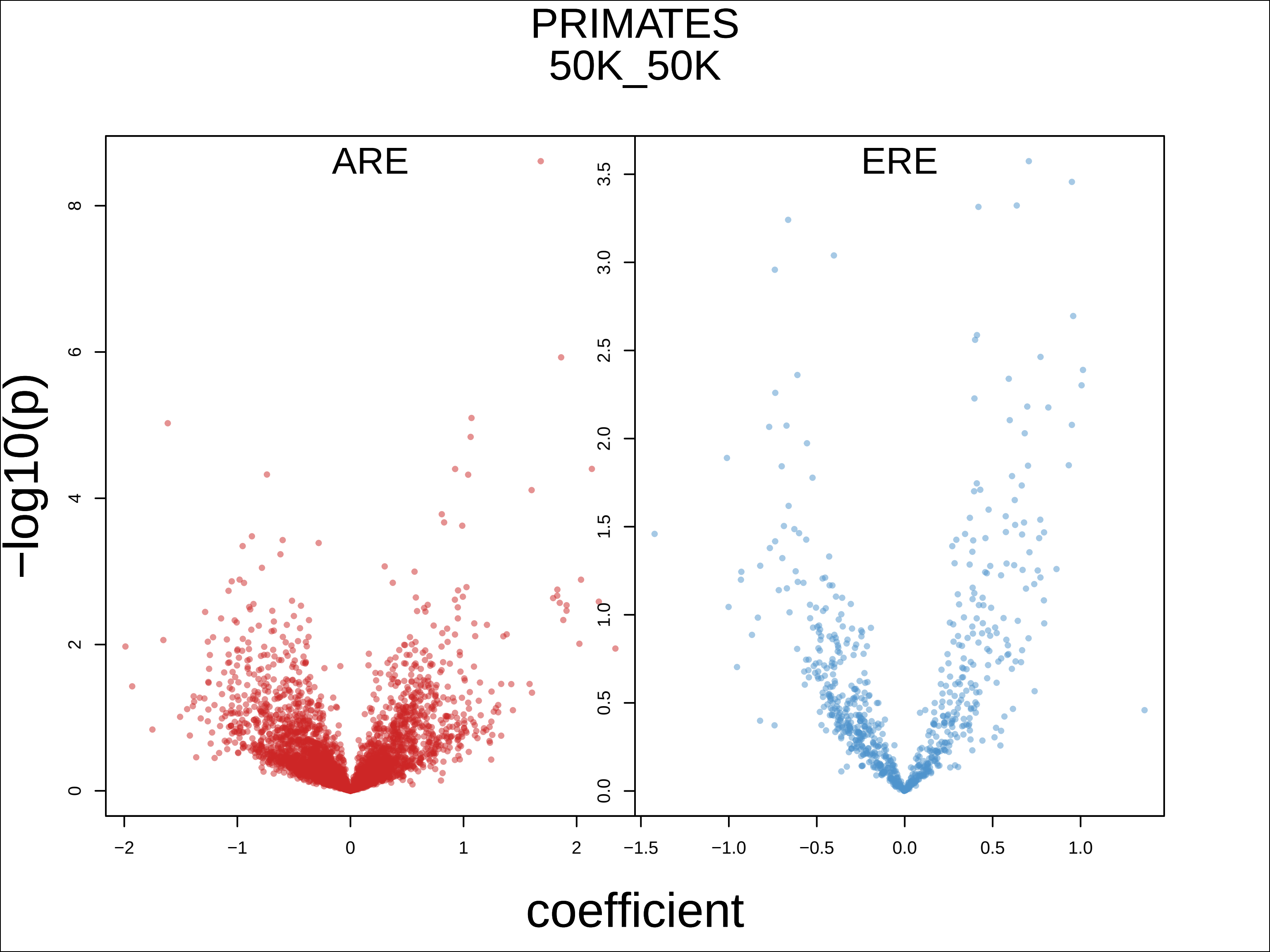
PRIMATES
50K_50K
ARE
ERE
−log10(p)
coefficient
0
2
4
6
8
0.0
0.5
1.0
1.5
2.0
2.5
3.0
3.5
−2
−1
0
1
2
−1.5
−1.0
−0.5
0.0
0.5
1.0

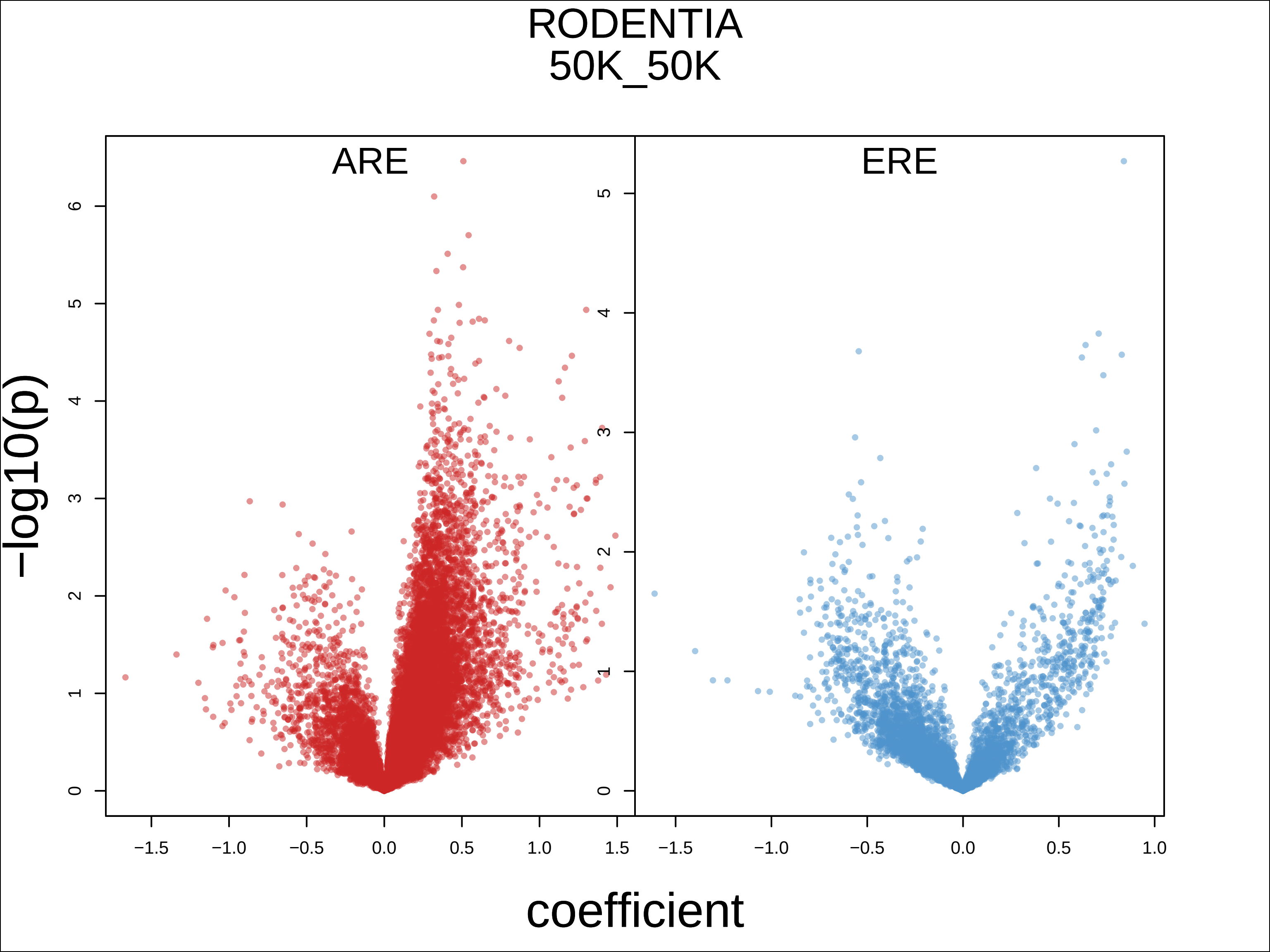

RODENTIA
50K_50K
ARE
ERE
−log10(p)
coefficient
0
1
2
3
4
5
6
−1.5
−1.0
−0.5
0.0
0.5
1.0
1.5

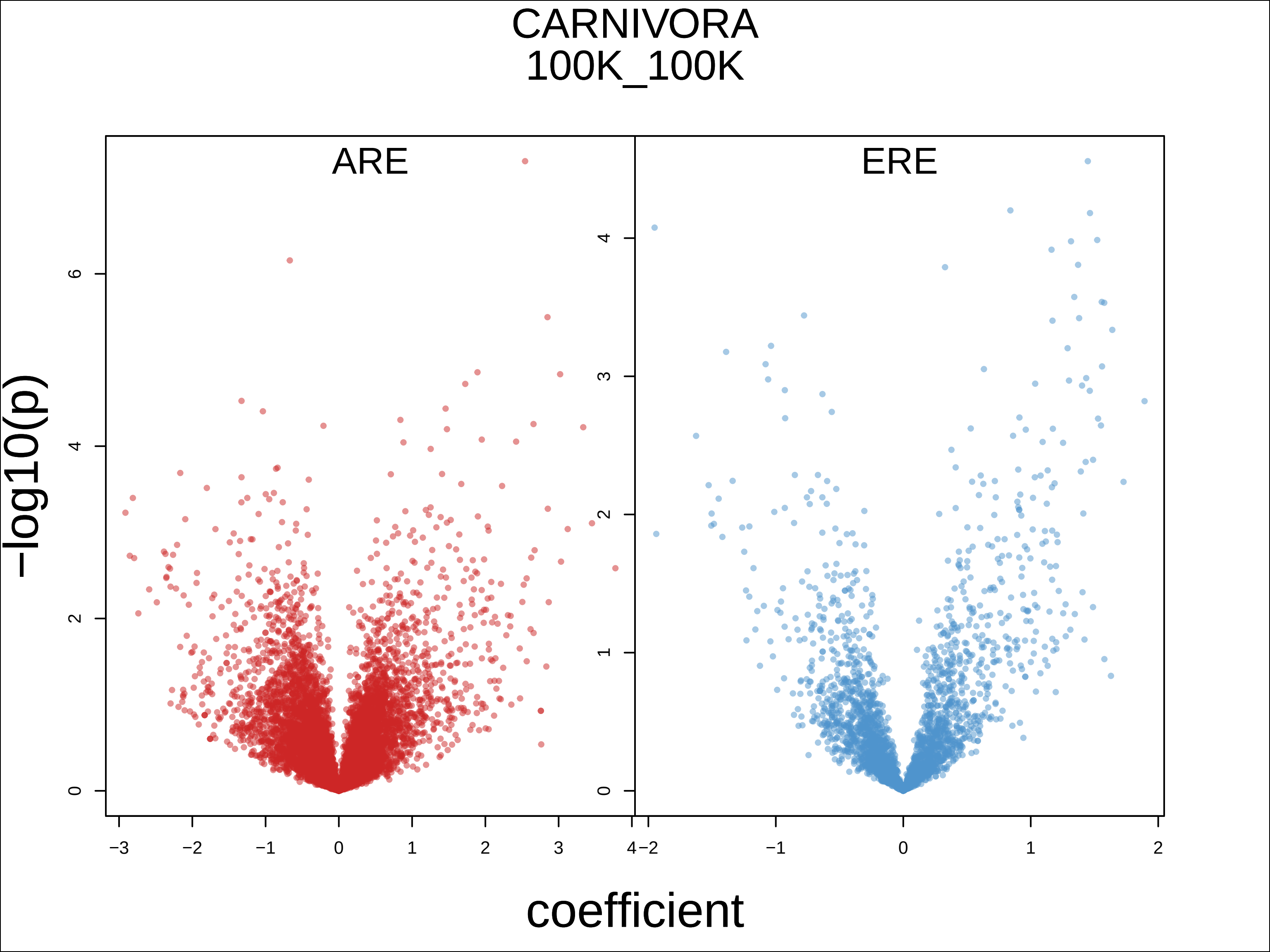
CARNIVORA
100K_100K
ARE
ERE
−log10(p)
coefficient
0
2
4
6
0
1
2
3
4
−3
−2
−1
0
1
2
3
4
−2
−1
0
1
2

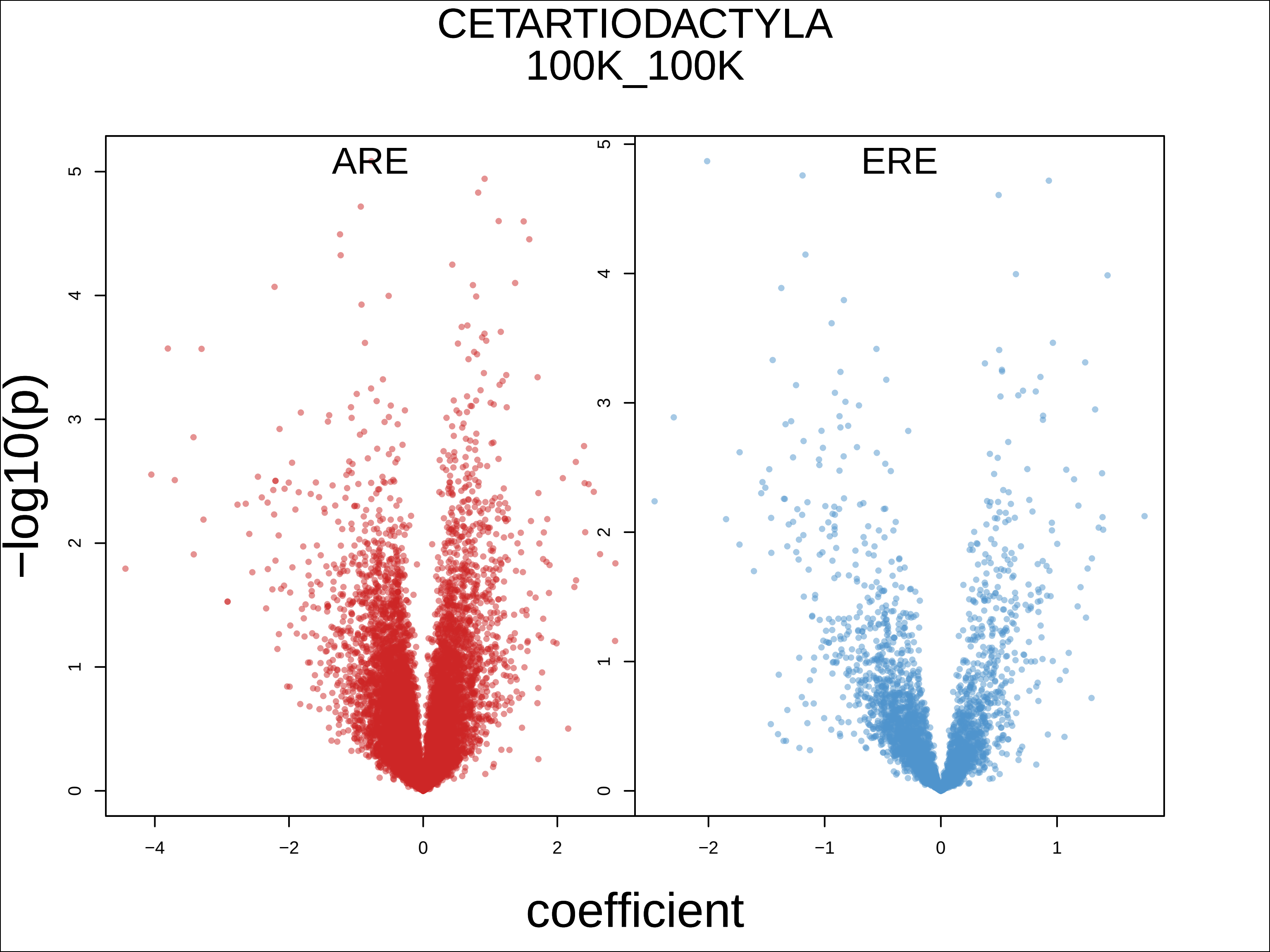

CETARTIODACTYLA
100K_100K
ARE
ERE
−log10(p)
coefficient
0
1
2
3
4
5
−4
−2
0
2
−1
1

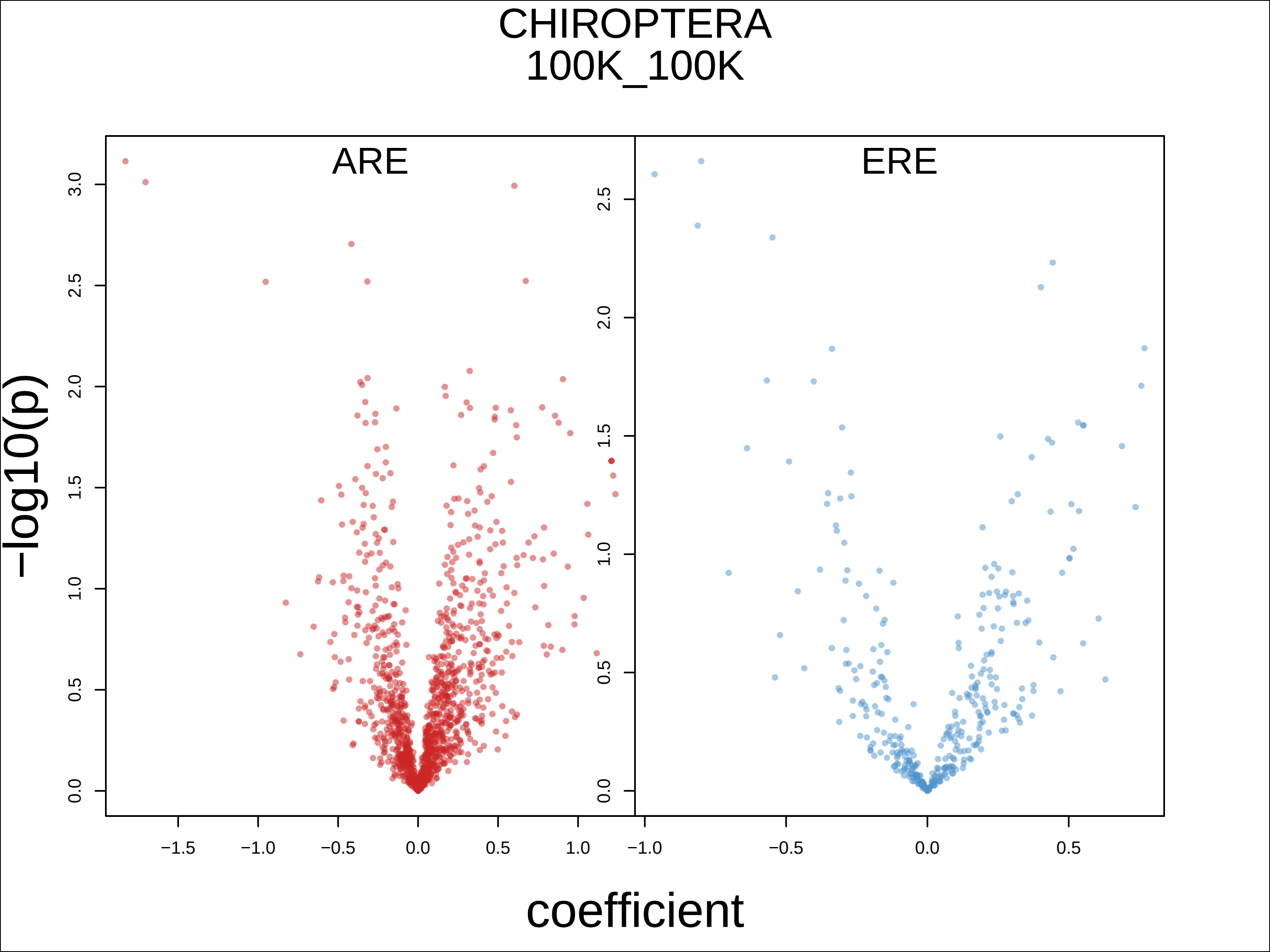

CHIROPTERA
100K_100K
ARE
ERE
-log10(p)
coefficient
3.0
2.5
2.0
1.5
1.0
0.5
0.0
−1.5
−1.0
−0.5
0.0
0.5
1.0

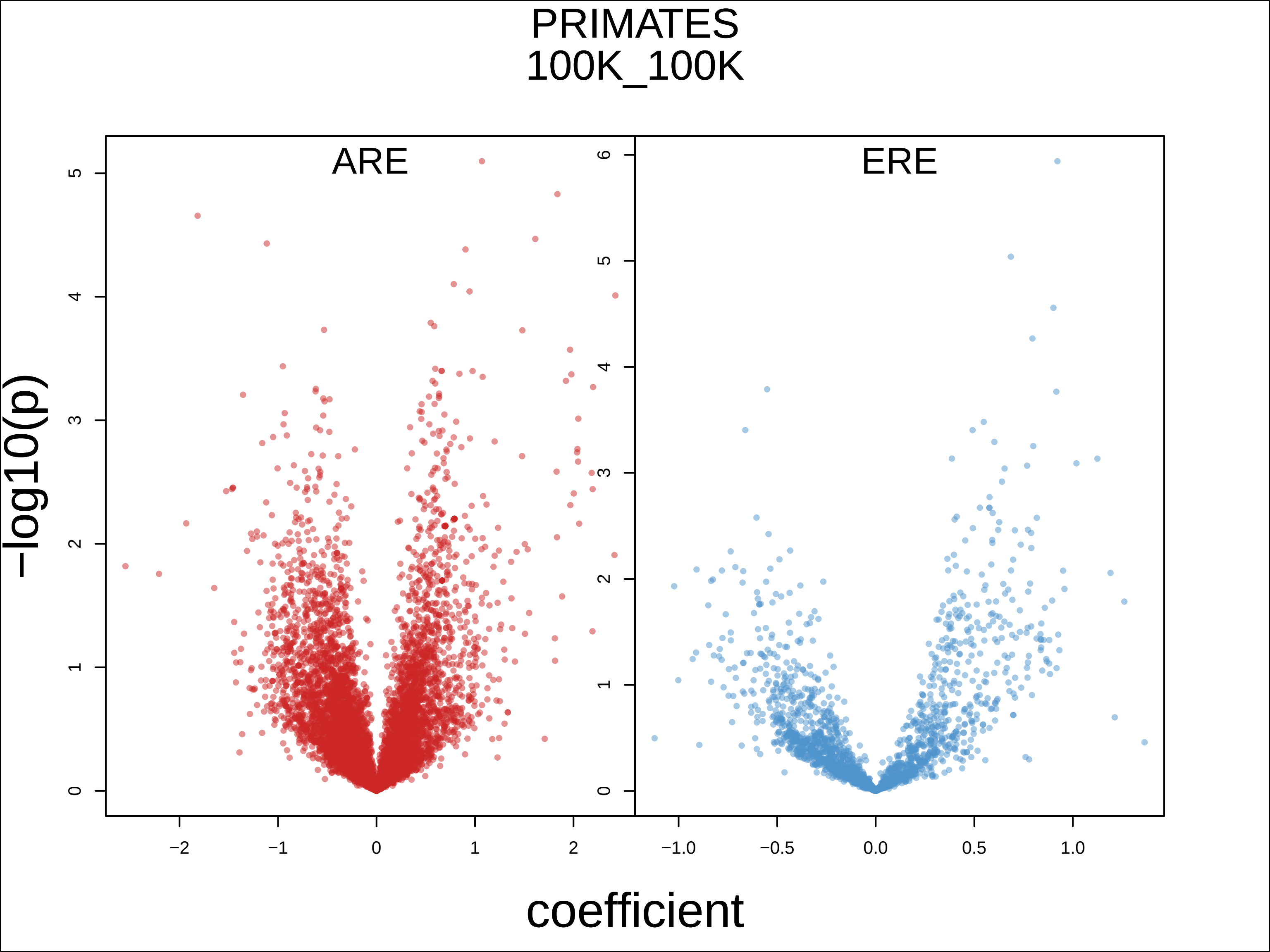
PRIMATES
100K_100K
ARE
ERE
−log10(p)
coefficient
0
1
2
3
4
5
6
−2
−1
0
1
2
−1.0
−0.5
0.0
0.5
1.0

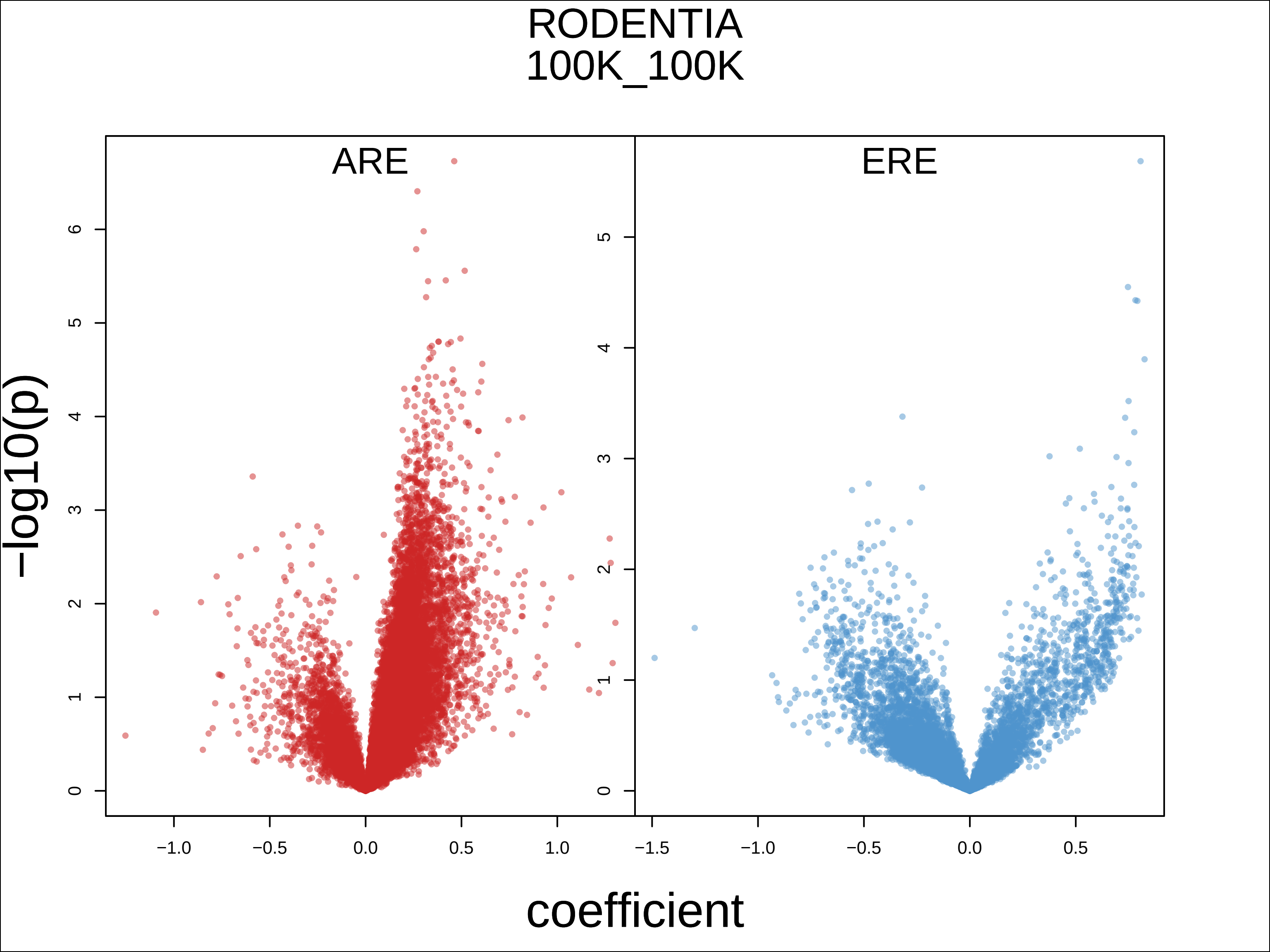
RODENTIA
100K_100K
ARE
ERE
−log10(p)
coefficient
0
1
2
3
4
5
6
−1.0
−0.5
0.0
0.5
1.0
−1.5

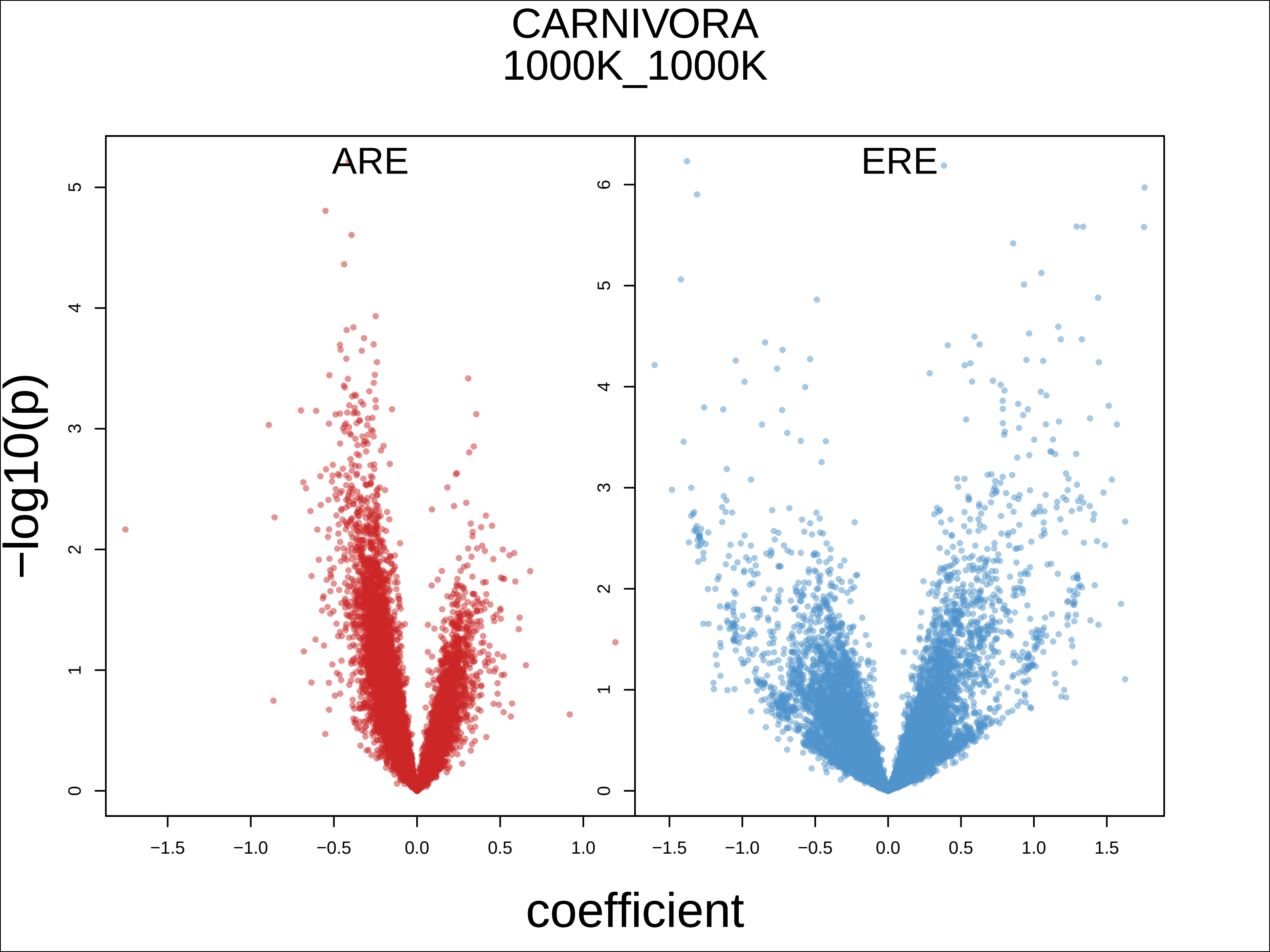
CARNIVORA
1000K_1000K
ARE
ERE
−log10(p)
coefficient
0
1
2
3
4
5
6
−1.5
−1.0
−0.5
0.0
0.5
1.0
1.5

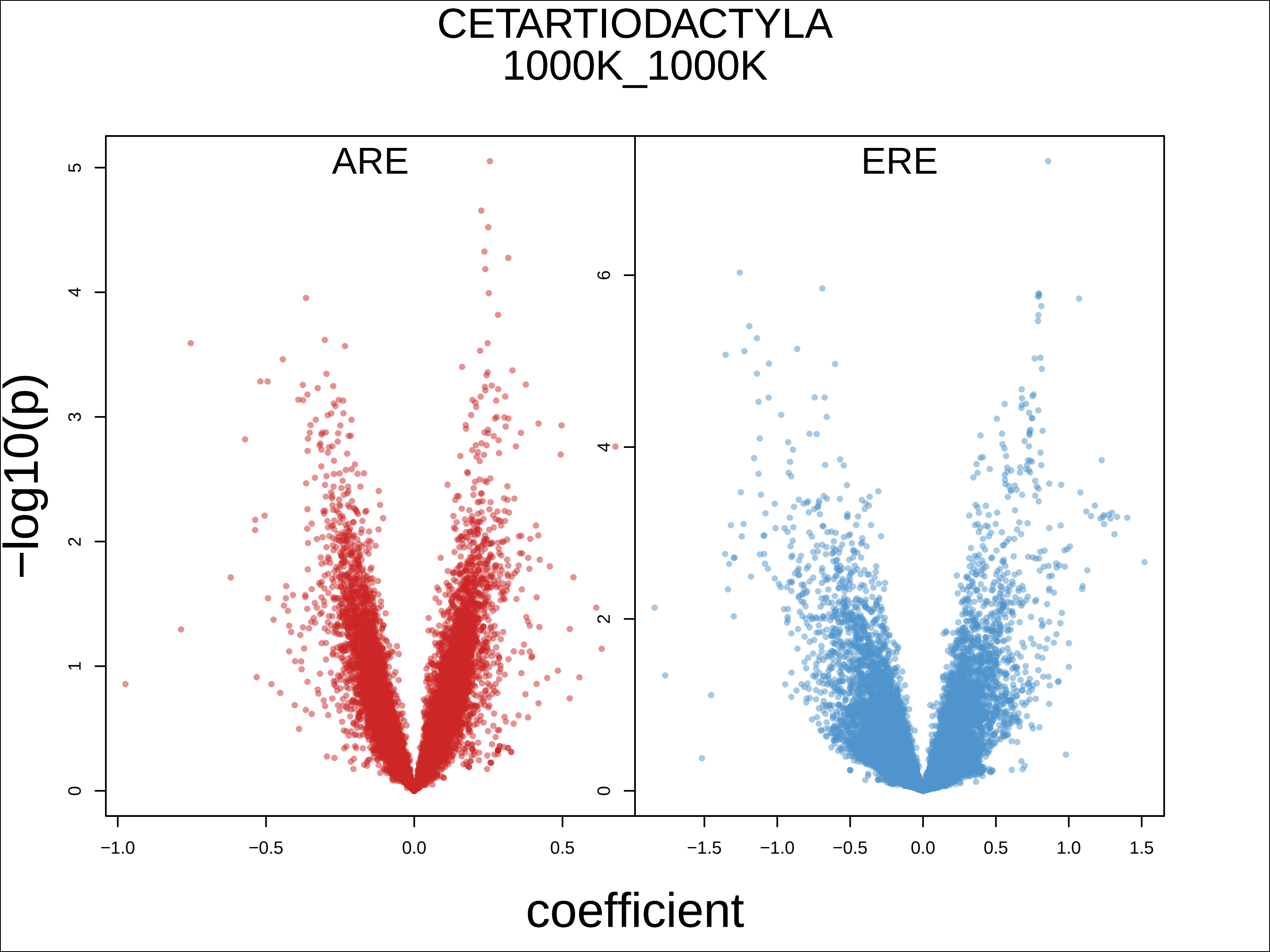
CETARTIODACTYLA
1000K_1000K
ARE
ERE
−log10(p)
coefficient
0
1
2
3
4
5
6
−1.0
−0.5
0.0
0.5
−1.5
1.0
1.5

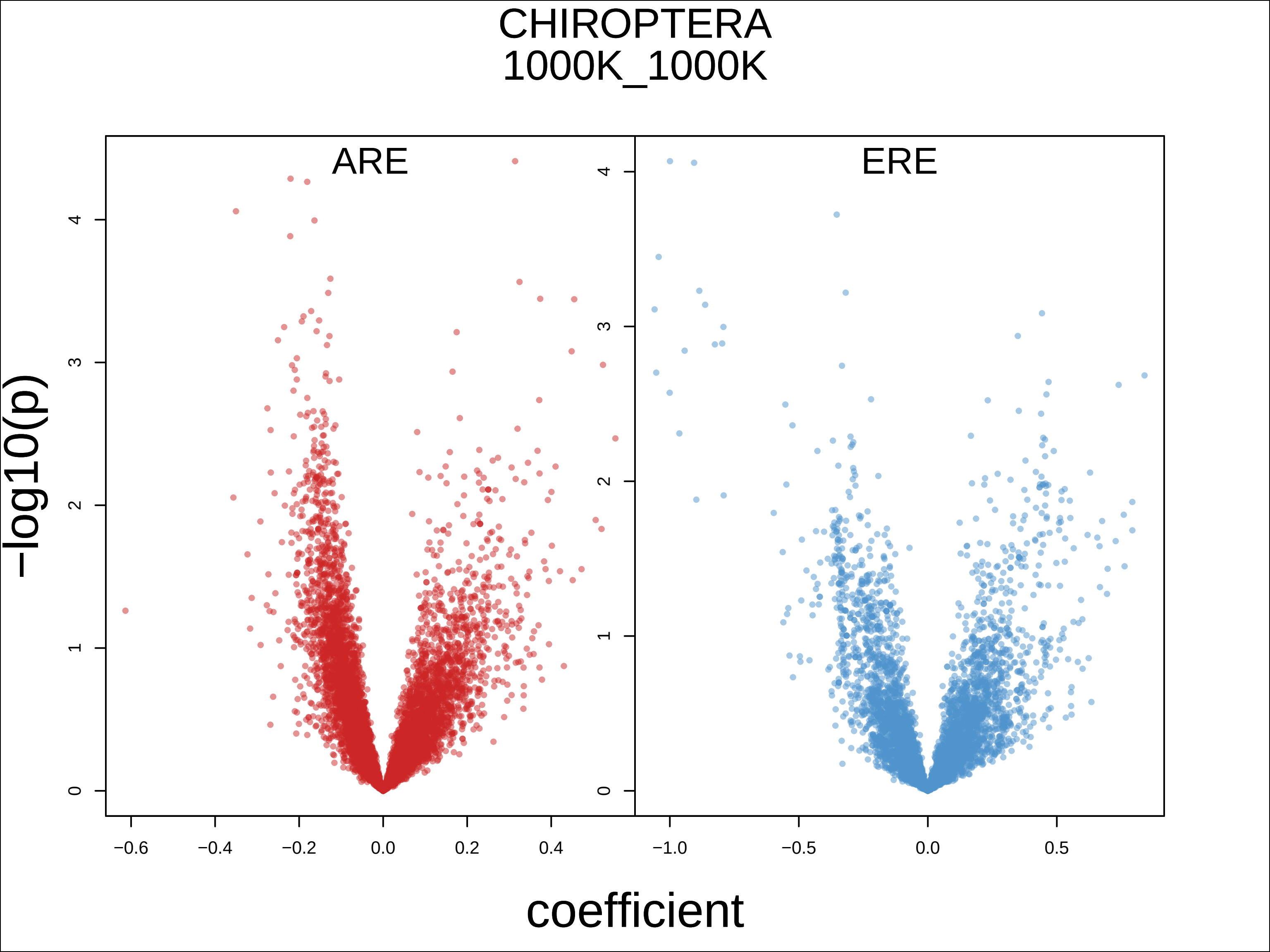

CHIROPTERA
1000K_1000K
ARE
ERE
−log10(p)
coefficient
0
1
2
3
4
−0.6
−0.4
−0.2
0.0
0.2
0.4
−1.0
−0.5
0.0
0.5

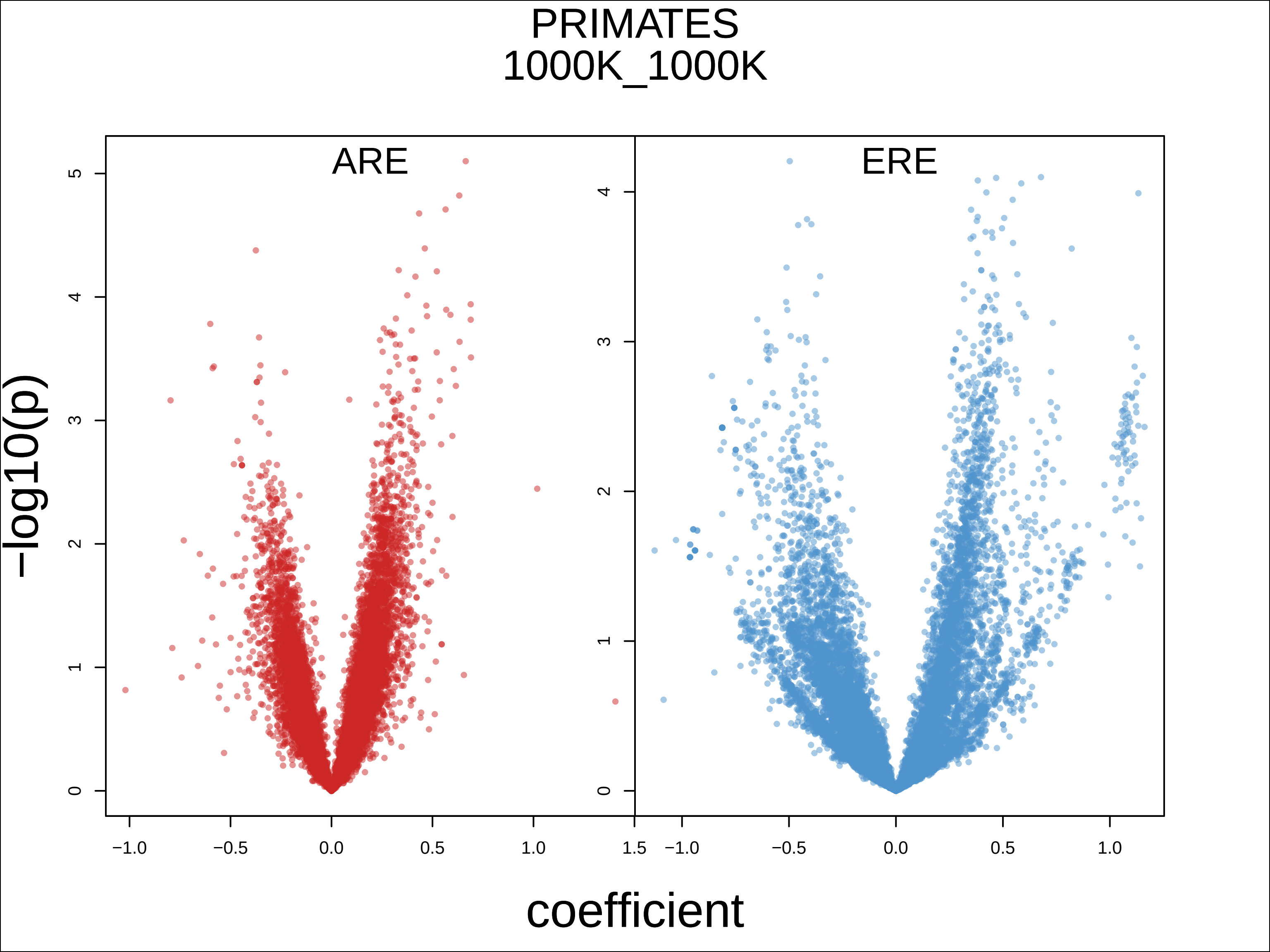

PRIMATES
1000K_1000K
ARE
ERE
−log10(p)
coefficient
0
1
2
3
4
5
−1.0
−0.5
0.0
0.5
1.0
1.5

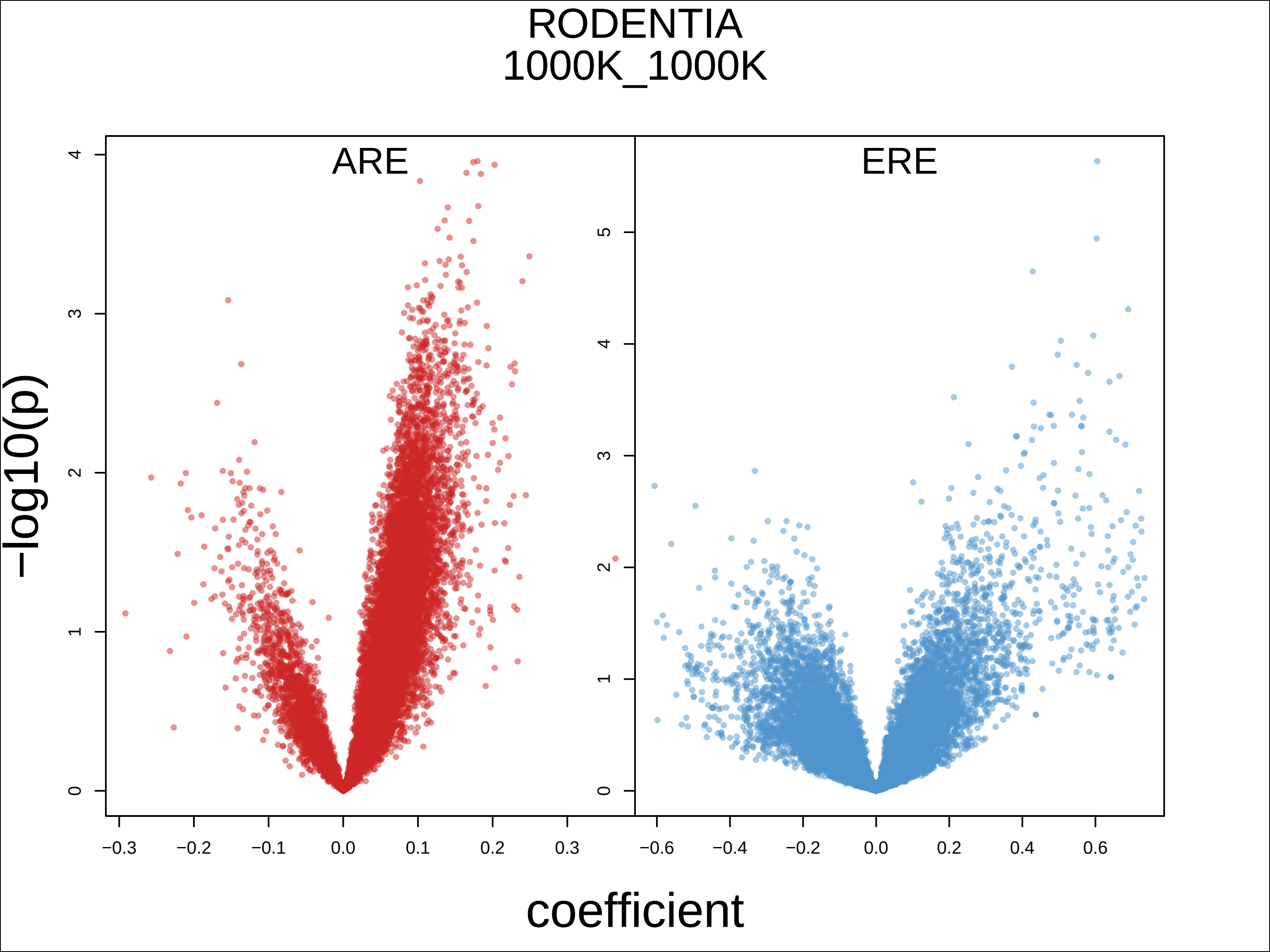

RODENTIA
1000K_1000K
ARE
ERE
−log10(p)
coefficient
0
1
2
3
4
5
−0.3
−0.2
−0.1
0.0
0.1
0.2
0.3
−0.6
−0.4
−0.2
0.0
0.2
0.4
0.6
